# Population genetics of *Paramecium* mitochondrial genomes: recombination, mutational spectrum, and efficacy of selection

**DOI:** 10.1101/281865

**Authors:** Parul Johri, Georgi K. Marinov, Thomas G. Doak, Michael Lynch

**Author notes:** Current address: Department of Genetics, Stanford University School of Medicine, Stanford, CA 94305, United States. These authors contributed equally to this work (Parul Johri, Georgi K. Marinov). **Corresponding author:** Parul Johri, Center for Evolution and Medicine, Life Sciences C, Room 360, 427 East Tyler Mall, Arizona State University, Tempe, AZ 85287.

## Abstract

The evolution of mitochondrial genomes and their population-genetic environment among unicellular eukaryotes are understudied. Ciliate mitochondrial genomes exhibit a unique combination of characteristics, including a linear organization and the presence of multiple genes with no known function or detectable homologs in other eukaryotes. Here we study the variation of ciliate mitochondrial genomes both within and across thirteen highly diverged *Paramecium* species, including multiple species from the *P. aurelia* species complex, with four outgroup species: *P. caudatum*, *P. multimicronucleatum*, and two strains that may represent novel related species. We observe extraordinary conservation of gene order and protein-coding content in *Paramecium* mitochondria across species. In contrast, significant differences are observed in tRNA content and copy number, which is highly conserved in species belonging to the *P. aurelia* complex but variable among and even within the other *Paramecium* species. There is an increase in GC content from ~20% to ~40% on the branch leading to the *P. aurelia* complex. Patterns of polymorphism in population-genomic data and mutation-accumulation experiments suggest that the increase in GC content is primarily due to changes in the mutation spectra in the *P. aurelia* species. Finally, we find no evidence of recombination in *Paramecium* mitochondria and find that the mitochondrial genome appears to experience either similar or stronger efficacy of purifying selection than the nucleus.

## INTRODUCTION

Mitochondrial genomes have played integral roles in furthering our understanding of relationships among species as well as revealing population structure and demographic history. As a consequence, we have obtained insights into the unique population-genetic properties of mitochondrial genomes. In most species, mitochondria are inherited uniparentally (but see Barr, et al. 2005). Although mitochondrial genomes are known to frequently undergo recombination in plants (Stadler and Delph 2002; Mackenzie 2007) as well as fungi (Fritsch, et al. 2014), no recombination has been detected in animals (Ballard and Whitlock 2004; Piganeau and Eyre-Walker 2004). Because of the unique mode of transmission and lack of recombination in some species, mitochondria have been suggested to have lower effective population sizes than their nuclear counterparts and therefore to accumulate more deleterious mutations (Lynch and Blanchard 1998; Neiman and Taylor 2009). In addition, mitochondrial genomes experience much higher spontaneous rates of mutation than their corresponding nuclear genomes in animals, but exhibit the opposite trend in plants (Lynch, et al. 2006).

Unlike the relatively uniform and conserved properties of metazoan mitochondrial genomes, mitochondrial genomes in unicellular eukaryotes exhibit remarkable variation in genome structure and GC content. Mitochondrial genome structures range from hundreds of short linear segments (0.3-8.3 kb) in the ichtysporean *Amoebidium parasiticum* (Burger, et al. 2003), an opisthokont, to many small (< 10 kb) circular genomes in the diplonemid *Diplonema papillatum* (Vlcek, et al. 2011), an excavate, to a single larger linear or circular chromosome, and a variety of other states (reviewed in Smith and Keeling 2015). In addition to variation in organization, mitochondria from unicellular lineages display widely diverse GC compositions, ranging from as low as 10% in some yeast (Smith 2012) to as high as 60% in *Lobochlamys culleus* (Borza, et al. 2009), although most species are AT rich, with an average GC content of 35% (Smith 2012).

Much has been learned about the structure, evolution and population-genetic environment of mitochondria in the main model systems, especially in plants and metazoans (Lynch 2007; Smith 2016). However, we lack such understanding of mitochondria of the majority of unicellular eukaryotes, where the bulk of eukaryotic phylogenetic diversity lies. We address this gap by surveying both within and between-species variation in mitochondrial genomes among multiple ciliate species belonging to the genus *Paramecium*.

In the large and morphologically and ecologically diverse ciliate lineage, mitochondrial genomes sampled so far are organized into large linear chromosomes, several tens of kilobases in length, with telomeres at the ends (Goddard and Cummings 1975; Morin and Cech 1988; Swart, et al. 2012). However, few mitochondrial genomes have until now been fully sequenced among the ciliates, with two in the *Paramecium* genus *(P. tetraurelia* and *P. caudatum*; (Barth and Berendonk 2011)), and only a few others: *Tetrahymena pyriformis* (Burger, et al. 2000), *Euplotes minuta* and *Euplotes crassus* (de Graaf, et al. 2009), *Oxytricha trifallax* (Swart, et al. 2012), *Stentor coeruleus* (Slabodnick, et al. 2017), *Ichthyophthirius multifiliis* (Coyne, et al. 2011), and the anaerobic ciliate *Nyctotherus ovalis* (de Graaf, et al. 2011).

While the *Paramecium* genus contains a number of distinct morphospecies, it is especially known for including a species complex consisting of multiple morphologically identical but sexually isolated species - the *P. aurelia* complex (Sonneborn 1975). Species in the *P. aurelia* complex are ancient, with the estimated time of divergence for the complex as a whole being on the order of 300 million years (McGrath, Gout, Johri, et al. 2014), implying that the genus *Paramecium* itself is even more ancient. Interestingly, among *Paramecium* species, there is an increase of GC content in mtDNA in the branch leading to the *P. aurelia* complex (Burger, et al. 2000; Barth and Berendonk 2011), allowing us to study the evolution of nucleotide composition across mitochondrial genomes that are structurally very similar.

The *Paramecium* species offer a particularly interesting system in which to study the evolution of mitochondrial genomes because of the unique population-genetic environment experienced by their cellular organelles. *Paramecium* cells are mitochondria-rich: each individual cell in *P. aurelia* species is estimated to contain about 5000 mitochondria, with about 8-10 genomes per mitochondrion (Beale and Tait 1981), which is much larger than in mammalian cells with 1000-2000 mitochondria (Kukat, et al. 2011) and yeast cells with 20-30 mitochondria per cell (Visser, et al. 1995). In addition, *Paramecium* lineages, like other ciliate species, possess two nuclei: the polyploid somatic nucleus (called the macronucleus), which divides amitotically where the bulk of transcriptional activity occurs, and the diploid germline nucleus (known as the micronucleus), which is transcriptionally silent, and which undergoes sexual reproduction. All *Paramecium* species can proliferate asexually for a limited number of generations, after which they senesce unless they undergo sexual reproduction or conjugation. During asexual proliferation, *Paramecium* undergoes binary fission during which mitochondria appear to double in length, replicate their genomes (Perasso and Beisson 1978), and are presumed to be randomly distributed between the two daughter cells. It therefore appears that mitochondria do not experience any bottleneck during mitotic division.

During conjugation, *Paramecium* cells exhibit cytoplasmic inheritance (Koizumi and Kobayashi 1989), i.e., despite the exchange of micronuclei between the two conjugants there is almost no exchange of cytoplasm and other organelles (reviewed in Meyer and Garnier 2002). Thus, mitochondria are uniparentally inherited. A distinct aspect of *Paramecium* mitochondrial biology is that *Paramecium* mitochondria appear to exist as independent structural units and do not undergo fusion, unlike the constant flux of organelle fusion and fission in other metazoan mitochondrial populations (Kiefel, et al. 2006). Both uniparental inheritance of mitochondria and the absence of fusion in the cytoplasm suggest a lack of recombination among mitochondria genomes.

In this study, we further the understanding of the biology and the population-genetic environment of ciliate mitochondria by presenting the complete mitochondrial genomes of nine species belonging to the *P. aurelia* complex, four outgroup (relative to *P. aurelia*) species, and additional 5-10 isolates for each of four *Paramecium* species. Using phylogenetic and population-genetic analyses, we investigate variation in protein-coding genes, tRNA content, and the forces governing the evolution of nucleotide composition of mitochondrial genomes across the phylogeny. Finally, we estimate the recombination rate across the genome and address the controversy of whether mitochondrial genomes experience reduced purifying selection in comparison to their nuclear counterparts.

## RESULTS

### Mitochondrial genome sequencing and assembly

We assembled complete mitochondrial genomes of seven species belonging to the *P. aurelia* complex: *P. biaurelia*, *P. tetraurelia*, *P. sexaurelia*, *P. octaurelia*, *P. novaurelia*, *P. decaurelia*, *P. dodecaurelia*, *P. quadecaurelia*, and *P. jenningsi*. In addition, we analyzed previously reported complete mitochondrial genomes of *P. tetraurelia and P. sexaurelia* belonging to the *P. aurelia complex*, as well as four outgroup species: *P. caudatum, P. caudatum-C026, P. multimicronucleatum*, and *P. multimicronucleatum-Peniche3I* (Figure 1; isolates sequenced in Johri et al., 2017). We note that *P. caudatum-C026* and *P. multimicronucleatum-Peniche3I* were initially sampled as individual isolates belonging to the *P. caudatum* and *P. multimicronucleatum* species (based on morphological criteria), respectively. However, the analysis of the mitochondrial sequences revealed that they are highly diverged from the reference strains (see below) and are therefore almost certainly separate species and were treated as such in subsequent analyses. For all 7 new mitochondrial genomes sequenced and assembled in this study, Illumina reads were assembled using SPAdes (Bankevich, et al. 2012), and mitochondrial contigs were identified by BLAST searches against the publicly available *P. caudatum* and *P. tetraurelia* mitochondrial sequences (see the Materials and Methods section for more details).

**Figure 1:**
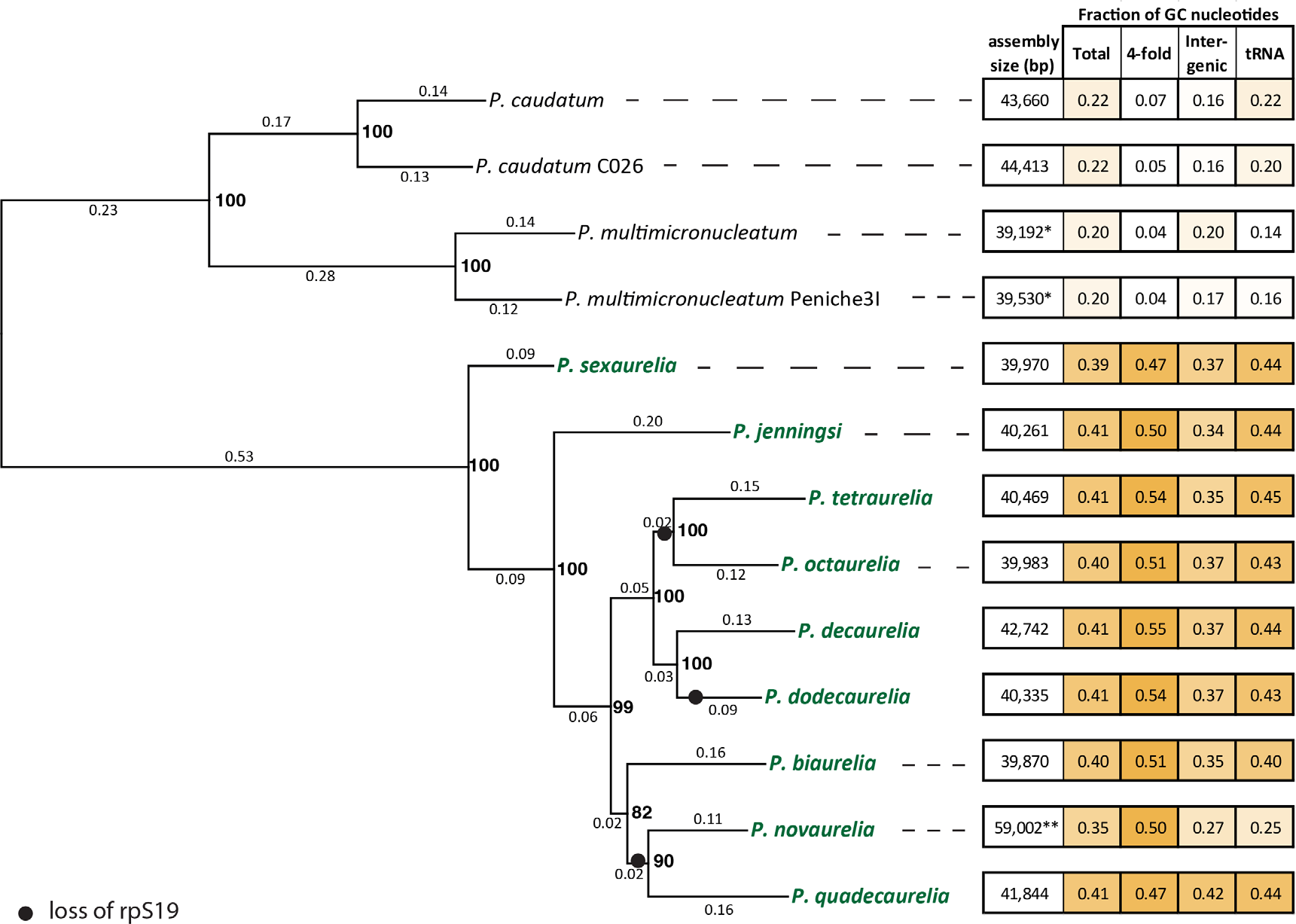
Mitochondrial phylogeny and change in nucleotide composition across *Paramecium* species. The species in green belong to the *P. aurelia* complex. Numbers on branches indicate total number of substitutions per site and bold numbers on nodes show bootstrap values. Black solid circles show the inferred event of loss of the ribosomal protein *rpS19*. The numbers on the right show the total assembly size (in bp) and mean GC content of the whole mitochondrial genome (Total), at 4-fold degenerate sites, at intergenic regions, and of tRNAs respectively. Starred (*) numbers indicate incomplete assembly of mitochondrial genomes.

In addition, we examined sequenced mitochondrial genomes of 10 isolates of both *P. tetraurelia* and *P. sexaurelia*, and 5 isolates of each *P. caudatum* and *P. multimicronucleatum* (Supplementary Figure 1) sampled worldwide (Johri, et al. 2017). Illumina paired-end reads from these isolates were mapped to the assembled reference genomes and SNPs were called as in Johri, et al. (2017). In this study, all mitochondrial genomes of individual isolates were also assembled *de novo* (Supplementary Figures 2, 3 and 4) in order to examine large-scale genome organization mapping.

### Genome structure, organization and telomeric repeats

All *Paramecium* mitochondrial genomes are linear (Goddard and Cummings 1975; Morin and Cech 1988; Swart, et al. 2012). Our assemblies show that they are ~40 kb in the *P. aurelia* species and ~44 kb in *P. caudatum* and *P. caudatum-C026*. Telomeric repeats and gene content observed at ends of the assembled genomes imply that the full lengths of the linear contigs have been assembled in most species with the exception of the *P. multimicronucleatum* and *P. multimicronucleatum* Peniche3I mitocontigs, which appear to be missing small portions of the 3′ end of the chromosome (Figure 2). The length of the assembled contigs in *P. multimicronucleatum* and *P. multimicronucleatum-Peniche3I* suggests an overall size closer to that observed in *P. aurelia* than to the larger mitochondrial genomes in the *P. caudatum* lineage. We also note that the raw assemblies for two of the *P. aurelia* species contain extensions (Supplementary Figure 3, 5, and 6). In *P. novaurelia*, an additional ~18 kb is present at the 5′ end of the mitocontig, while in *P. quadecaurelia*, a small, ~1 kb extension is seen at the 3′ end. However, the read coverage over these regions is very different from the rest of the mitocontigs, suggesting either misassembly or heterogeneity within cell populations. We therefore ignored these extensions in subsequent analysis. The organization of the genomes is very similar, with gene order preserved almost perfectly between all species (Figure 2).

**Figure 2:**
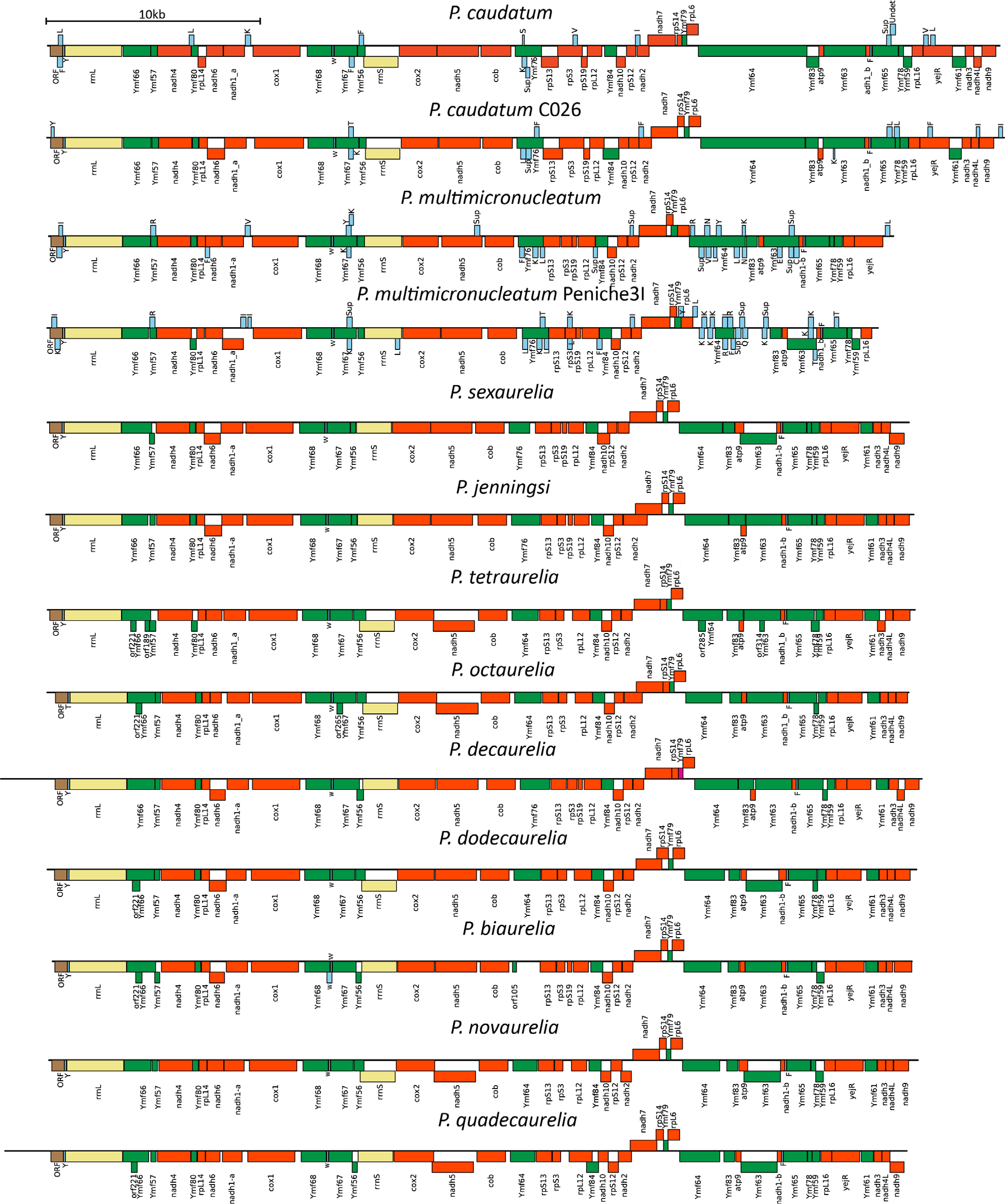
Structure of mitochondrial genomes in *Paramecium*. All *Paramecium* mitochondrial genomes are linear with 24 protein-coding genes (shown in reddish orange), 2 rRNA genes (yellow), lineage-specific genes referred to as *Ymf* genes (in green) and varying numbers and content of tRNA genes (shown in light blue).

We also identified telomeric repeats based on the sequences at the end of assembled mitocontigs (see the Methods section for more details). In most *P. aurelia* species, we identify almost identical repeats, with a 23-bp consensus sequence GCCCTGGTGGCCCTAGAAGCTCC (Figure 3). However, in *P. jenningsi* and *P. sexaurelia* the telomere repeat motif is the same length, but has 1 nucleotide difference from the consensus sequence. Of note, these two species are the earliest diverging ones within the *P. aurelia* complex species included in our analysis. We observed even more divergent telomeric sequences in *P. caudatum* (GCCCTGGTAACGCTGGTCGCCCTTTTAAAATA) and *P. multimicronucleatum* (GCCCTTGTTACACTTGGTGGCTCTTTAAGCTCT). In these species, the core telomeric repeat sequence has been expanded by an additional 10-bp of sequence not present in *P. aurelia*. The *Paramecium* core telomeric repeat is broadly similar to that of *Tetrahymena* (ACCCTCGTGTCTCTTTA; Figure 3), the other Oligohymenophorean genus for which mitochondrial genomes are available, but distinct from what is observed in distantly branching ciliates such as the spirotrichean *Oxytricha trifallax* (Figure 3).

**Figure 3:**
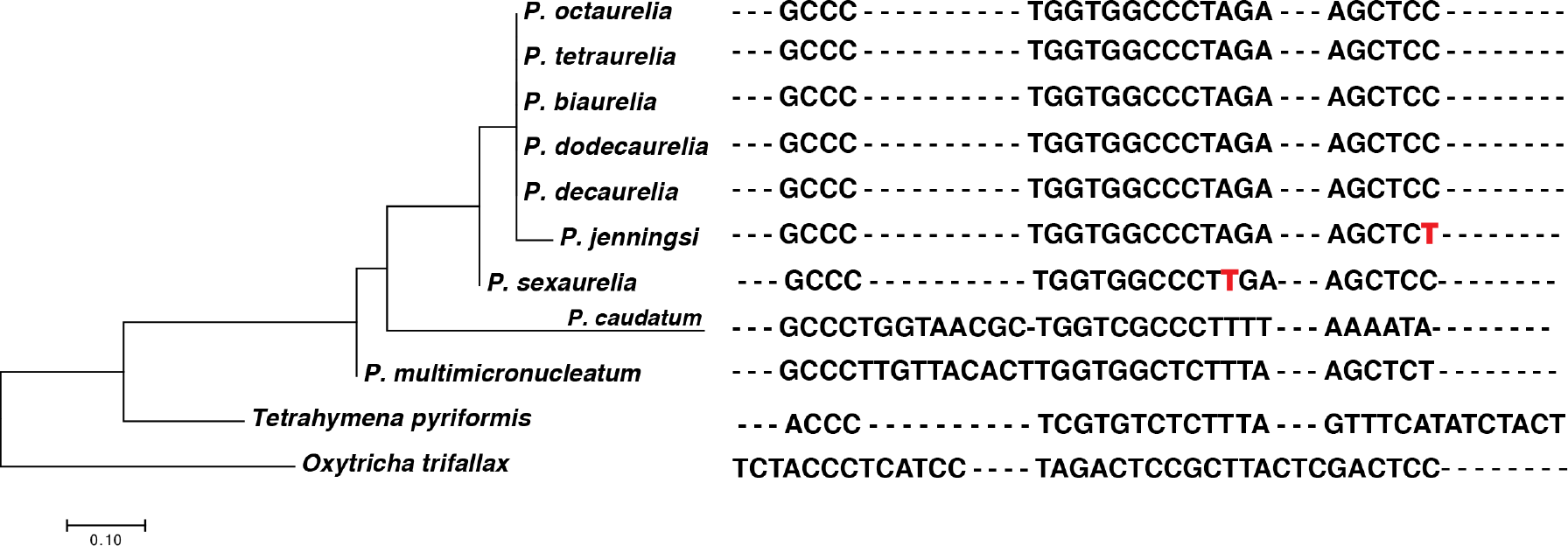
Telomeric repeat sequences in the *Paramecium* genus. Nucleotides in red show the single base pair differences among the *P. aurelia* species. The phylogenetic tree on the left is built using the telomeric repeat sequences.

**Figure 4:**
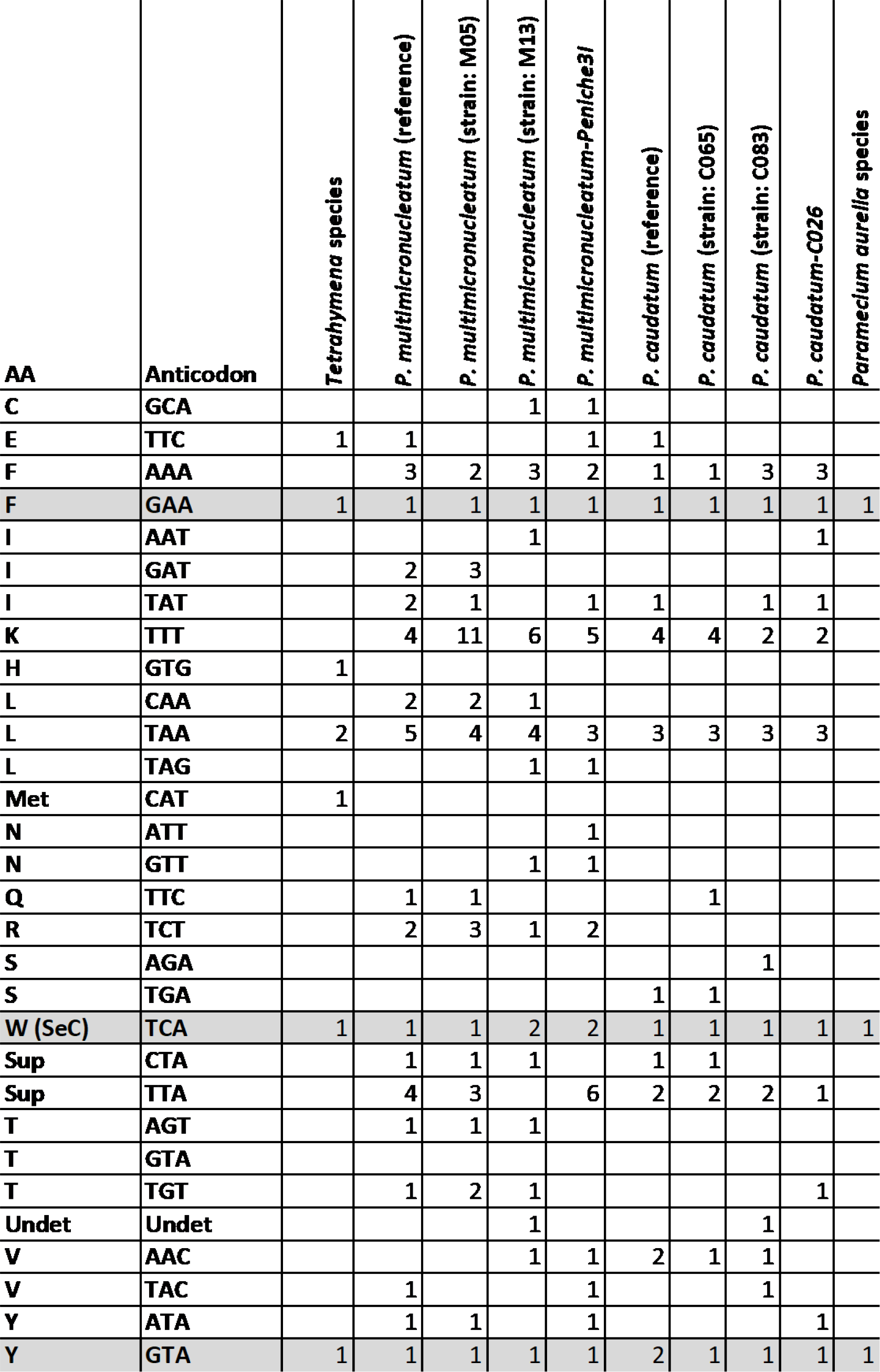
tRNA content variation in *P. caudatum* and *P. multimicronucleatum* strains. tRNAs shaded in grey are present in all *Paramecium* and *Tetrahymena* species. tRNAs whose anticodons were ambiguous are displayed as “Undet”.

Examination of existing RNA-seq datasets for several of the species revealed no evidence for RNA editing in *Paramecium* mitochondria (data not shown), in concordance with previous reports (Orr, et al. 1997).

### Variation in protein coding gene content between and within species

All *Paramecium* mitochondrial genomes contain a core set of 15 genes (Figure 2) involved in electron transport and ATP synthesis (*atp9*, *cob*, *cox1*, *cox2*, *nadh1*, *nadh2*, *nadh3*, *nadh4*, *nadh4L*, *nadh5*, *nadh6*, *nadh7*, nadh9 and *nadh10*), the heme maturase *YejR*, and 8 ribosomal protein genes (*rpL6*, *rpL12*, *rpL14*, *rpL16*, *rpS3*, *rpS12*, *rpS13*, and *rpS14*). One ribosomal protein subunit *rpS19* was found to be absent from *P. tetraurelia*, *P. octaurelia*, *P. novaurelia*, *P. dodecaurelia*, and *P. quadecaurelia* and therefore appears to have been independently lost from mitochondrial genomes at least 3 times along the *P. aurelia* phylogeny (Figure 1). Remarkably, we also find a presence-absence polymorphism of *rpS19* within *P. sexaurelia* isolates, with the ORF not being identifiable in 2 out of 11 isolates studied (Supplementary Figure 7). Although this might suggest a mitochondrial gene in the process of transfer to the nucleus, the closest hit in the nuclear genome has an E value of 0.27, implying either a complete loss or perhaps transfer to a mitochondrial plasmid, which has not been captured in existing assemblies.

*Paramecium* genomes also contain 16 ciliate-specific genes of unknown function, called *Ymf* genes (Burger, et al. 2000), named according to the Commission on Plant Gene Nomenclature (Price and Reardon 2001). Each sequenced strain contains all of these genes, with the exception of *P. multimicronucleatum* strain M13 in which *Ymf80* and *Ymf83* appear to have fused with *nadh4* and *Ymf64*, respectively. In addition, several ORFs originally identified in *P. tetraurelia*, are observed in only one or a few additional *P. aurelia* mitochondrial genomes, typically overlapping longer ORFs: *orf189, orf285*, and *orf314* restricted to the three *P. tetraurelia* isolates; *orf221* present in *P. tetraurelia*, *P. octaurelia* and *P. biaurelia*; *orf265* in *P. biaurelia*; and *orf78* found only in *P. tetraurelia* isolate A.

In addition, we found a previously unidentified ORF larger than 100 amino acids, which appears to be present in all *Paramecium* mitochondrial genomes and is located at the very 5′ end of the chromosome, immediately before the large ribosomal RNA. This ORF has homology to the CRISPR-associated endoribonuclease Cas6; its functional significance is unclear at present. It could potentially represent a case of horizontal gene transfer from endosymbiotic bacteria (Preer 1969; Fokin and Gortz 2009) often associated with *Paramecium*. Sequencing of more ciliate mitochondrial genomes could provide insight into its origin.

### *Paramecium* species are highly diverged and contain cryptic species complexes

We found most species to be fairly evolutionarily distant from each other, as measured by the average number of nonsynonymous (*dN*) and synonymous (*dS*) site substitutions per site (Supplementary Figure 8) between all pairs of species, using yn00, PAML (Yang 2007; version 4.9a). *P. decaurelia* and *P. dodecaurelia* (average *dS*: 0.85; average *dN*: 0.05, average total divergence: 0.23) are the closest species pair, followed by *P. tetraurelia* and *P. octaurelia* (average *dS*: 0.93; average *dN*: 0.09; average total divergence: 0.27). All other species pairs exhibit on average *dS* > 1.0, i.e., synonymous sites in protein coding genes have on average undergone more than 1 substitution each since the time of divergence. We found that mean *dS* between *P. caudatum* and *P. caudatum-C026* is ~2.0 (average *dN*: 0.07; average total divergence: 0.28), implying that these two isolates possibly represent separate species. A similar observation confirms that *P. multimicronucleatum-Peniche3I* and *P. multimicronucleatum* are probably separate species (average *dS* = 1.97; average *dN*: 0.10; average total divergence: 0.26). Our observations suggest the possibility of cryptic species complexes in both *P. caudatum* and *P. multimicronucleatum*, as previously suspected (Hori, et al. 2006; Tarcz, et al. 2012). We also note that several additional *P. multimicronucleatum* isolates most closely related to *P. multimicronucleatum-Peniche3I* might also be separate species as they are fairly divergent from the reference strain as well as *P. multimicronucleatum-Peniche3I* (Supplementary Figure 1). The phenomenon of species complexes containing numerous morphologically identical species, already known for *P. aurelia* (Sonneborn 1937) and *T. pyriformis* (Gruchy 1955) is therefore probably much more widespread among ciliates than previously appreciated.

### Selective pressures on ciliate-specific mitochondrial genes

The 16 ciliate-specific mitochondrial genes found in all *Paramecium* species have orthologs in *Tetrahymena* species as well as in *Oxytricha*. Thus, these genes have been preserved for a long evolutionary time, and yet have diverged sufficiently that no known homologs exist in well-studied species in other eukaryotic kingdoms. Across the *Paramecium* species, they appear to be on average faster evolving (Supplementary Figure 9, Supplementary Table 1) with average *dN/dS* ≅ 0.11, in comparison to genes that encode enzyme components of the respiratory chains (average *dN/dS* ≅ 0.04) and ribosomal proteins (average *dN/dS* ≅ 0.06).

The *dN*/*dS* values here were obtained for each gene under the assumption that *dN*/*dS* remains constant across the phylogeny using CODEML, PAML (Yang 2007; version 4.9a). Ciliate-specific *Ymf* genes therefore exhibit a relatively higher rate of evolution than other genes, with as little as 30% sequence identity between *P. aurelia*, *P. caudatum*, and *P. multimicronucleatum*, similar to observations in *Tetrahymena* (Moradian, et al. 2007), as well as with higher values of π_n_/π_s_ in *P. tetraurelia*, *P. sexaurelia*, *P. caudatum*, and *P. multimicronucleatum* (Supplementary Figure 10).

The higher rate of evolution of *Ymf* genes can either be explained by relaxed purifying selection at the sequence level, or recurrent positive selection over long periods of time, or a combination of both. To distinguish between these possibilities, we performed a McDonald-Kreitman test (MK test) in the species *P. tetraurelia*, *P. sexaurelia*, *P. caudatum*, and *P. multimicronucleatum*. As most of these species are highly diverged from each other, we used ancestral reconstruction over the set of all 13 taxa to first infer ancestral nucleotides for each internal node. The numbers of synonymous (*D*_*S*_) and nonsynonymous (*D*_*N*_) changes were then inferred along each terminal branch leading to each of the 4 species mentioned above. We tested for positive selection by determining whether the proportion of nonsynonymous fixed differences relative to synonymous ones was significantly larger than the proportion of nonsynonymous relative to synonymous polymorphic differences. None of the genes in *P. tetraurelia* were found to have undergone positive selection by this criterion (Holm–Bonferroni corrected *p* ≥ 0.05 in each case). In *P. sexaurelia*, *cox1*, *Ymf76*, and *Ymf63* are the only genes that showed significant positive selection, whereas in *P. caudatum*, only *Ymf64* exhibits positive selection (Supplementary Table 2). With very few SNPs available in *P. multimicronucleatum*, the MK test was not significant for any gene. Nevertheless, we find no statistical enrichment for number of genes evolving under positive selection among the *Ymf* relative to other genes in *P. sexaurelia* (*p* = 0.56; Fisher’s exact test) and *P. caudatum* (*p* = 0.40; Fisher’s exact test), suggesting that the majority of them are likely evolving fast due to relaxed sequence constraints.

It is possible that most of the *Ymf* genes are simply highly diverged genes that perform the same functions as genes found in other mitochondrial genomes, but are not identifiable by sequence. In order to gain some insight into their possible functions, we used HHPred (Soding, et al. 2005) to predict their homologs and found that parts of *Ymf* genes display homology to standard mitochondrial genes, encoding ribosomal subunits and proteins involved in electron transport. About 37% of sequence from *Ymf59* was found to be homologous to *rpS10*, ~22% sequence of *Ymf61* to a 30S ribosomal protein S24E, ~28% of *Ymf64* sequence to a 30 S *rpS3*, ~79% of *Ymf65* sequence to the NADH subunit and ~11% of *Ymf67* to cytochrome c oxidase, subunit *P. Ymf59* and *Ymf64* in *Oxytricha* were also previously found to be homologous to *rpsS10* and *rpS3* respectively (Swart, et al. 2012).

### tRNA content variation across and within species

While the number and identity of protein-coding genes is generally the same across the genus, tRNAs display significant variation. All species in the *P. aurelia* complex have only three tRNA loci, at highly conserved locations (Figure 2), most of the time consisting of a Y, F, and W tRNAs (the latter recognizing the UGA stop codon that has been reassigned to tryptophan in ciliate mitochondria). The one exception is *P. octaurelia*, in which a T tRNA has been substituted for the Y tRNA that other species possess. In stark contrast, large and highly variable sets of tRNA are predicted in mitochondrial genomes in the *P. caudatum* and *P. multimicronucleatum* lineages (Figure 3), with as many as 38 tRNA predictions in *P. multimicronucleatum* M05, and with individual *P. caudatum* and *P. multimicronucleatum* isolates having different tRNAs at multiple loci in their mitochondrial genomes. These tRNAs often overlap protein-coding genes (in contrast to what is observed in *P. aurelia*), in particular *Ymf* genes. Of note, the three tRNAs found in the *P. aurelia* species are also present in all outgroup species. The absolute numbers of the tRNAs found should be taken with caution, as some of the predicted tRNAs overlap each other as well as *Ymf64* in the *P. multimicronucleatum* strains. However, even if these are excluded, the number of tRNAs in both *P. caudatum* and *P. multimicronucleatum* strains is still large and variable and consist of tRNAs not found in the *P. aurelia* species. This is a unique finding among mitochondrial genomes and should be investigated more thoroughly in future studies to better understand the origin of the tRNAs.

### Mutation spectrum and GC-content variation within the *Paramecium* genus

Perhaps the most remarkable discontinuity in *Paramecium* mitochondrial genomes is the difference between the high GC content observed in species of the *P. aurelia* complex (~40%) and the low GC content in the outgroup species (~20%). Very low GC content is also observed in mitochondrial genomes of other ciliates, for instance in the five *Tetrahymena* species (~18-21%), in *Icthyophthirius multifiliis* (~16%) (Coyne, et al. 2011), and in *Oxytricha trifallax* (~24%) suggesting a lineage-specific increase in GC content along the branch leading to *P. aurelia* species. In contrast, nuclear genomes have relatively similar levels of GC content across all *Paramecium* species (McGrath, Gout, Doak, et al. 2014). The low GC content of *P. caudatum* and *P. multimicronucleatum* mtDNA is marked by a highly biased codon usage (Barth and Berendonk 2011), with most synonymous positions exhibiting a strong bias for A or T nucleotides (Supplementary Figure 11). Indeed, the difference in GC content is most pronounced at 4-fold degenerate sites, with the GC content at such sites being as low as 3.5% in *P. multimicronucleatum* and as high as 54.5% in *P. decaurelia* (Figure 1). The o-fold redundant sites in the outgroup species have much higher GC content (~26-37%) and the GC content (31-36%) of the rRNAs is similar across the entire genus suggesting strong selection on rRNA nucleotide content.

At least three forces may be responsible for determining the GC content at 4-fold degenerate sites: (1) mutational processes, (2) codon usage bias due to selection, and/or (3) genome-wide selection for higher/lower GC content. We therefore examined GC composition in other regions of the genomes, such as tRNAs and intergenic regions, which do not experience selection for codon usage. Both tRNAs and intergenic regions have elevated GC content in the *P. aurelia* species (~40-45% and ~34-37%) relative to *P. caudatum* and *P. multimicronucleatum* (~14-22% and ~16-20% respectively; Figure 1), suggesting that either differences in mutational bias or selection for genome-wide GC content is primarily responsible for changes in nucleotide composition across the genome.

To distinguish between mutation and selection as the possible explanations for these observations, we evaluated the mutational pressures acting on *Paramecium* species, using two approaches. First, we used the mitochondrial mutation spectrum obtained from mutation-accumulation (MA) studies of *P. tetraurelia* (Sung et al, 2012) to calculate the AT mutation bias, *m* = *v*/*u*, where *v* is the mutation rate from G/C to A/T and *u* is the rate of mutation from A/T to G/C (Table 1). We found m ≅ 1.0, showing little mutation bias and corresponding to expected equilibrium GC content of 51.3% (Table 1). This is very close to the GC content of 4-fold redundant sites (54%) in *P. tetraurelia*. We also analyzed existing sequencing data from MA experiments in *P. biaurelia* and *P. sexaurelia* (Long, Doak, et al. 2018; Supplementary Table 3). These data imply slight mutation bias towards GC in *P. biaurelia* and *P. sexaurelia*, corresponding to expected equilibrium GC percentages (Sueoka 1993) of 55.8% and 67.0% respectively. Given the relatively small number of mutations (a few tens) identified in each set of MA experiments, there is considerable uncertainty in these estimates. Nonetheless, these data do not present evidence that 4-fold degenerate sites in *P. aurelia* mitochondria are evolving away from mutation equilibrium via selection.

**Table 1:**
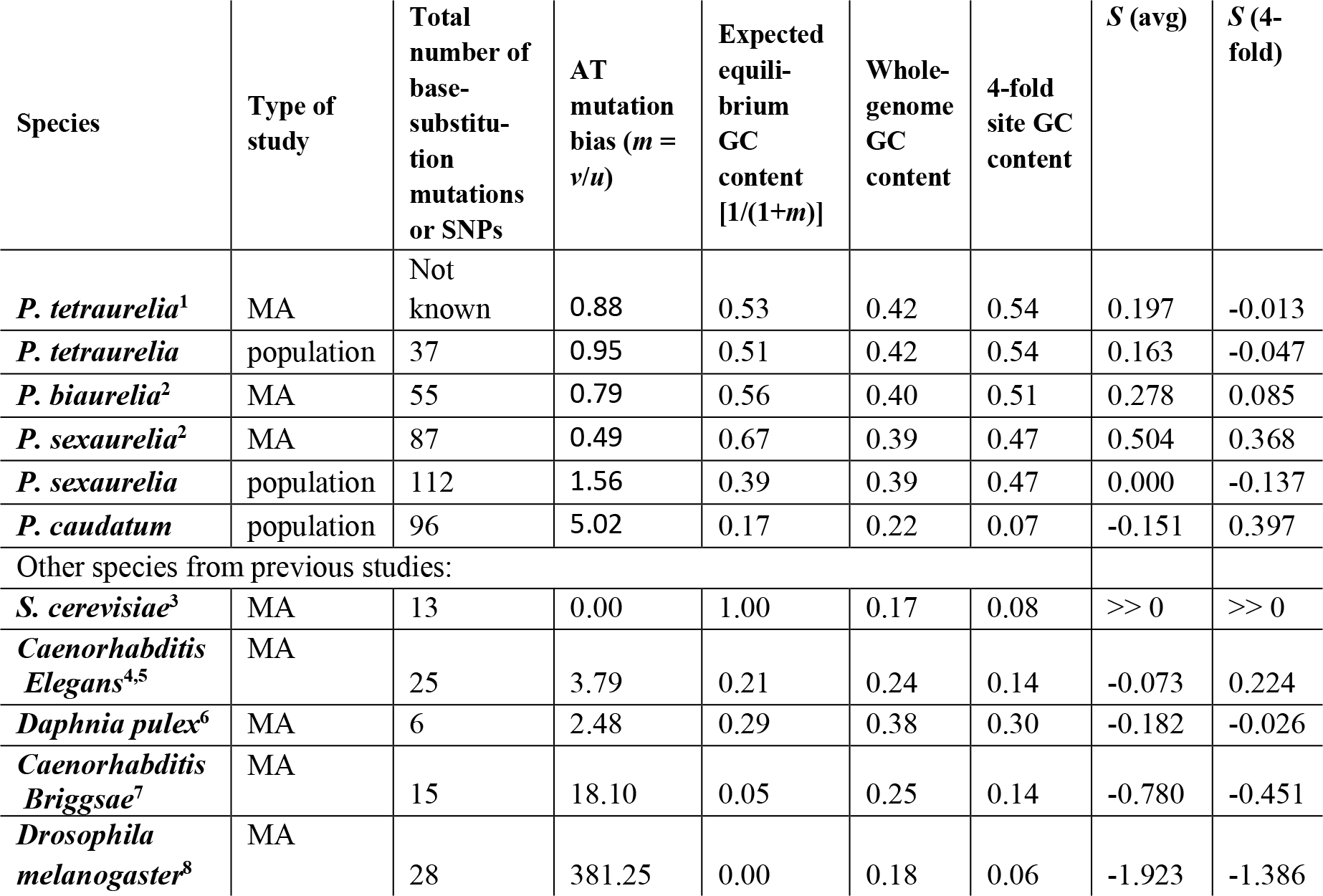
Mutation bias towards AT (*m*) in the mitochondrial genomes of *Paramecium* and other species, where *v* and *u* are mutation rates from G/C to A/T, and A/T to G/C respectively. Values of *m* less than 1 indicate that mutation spectrum is biased towards G/C. *S* = 4*N*_e_*S* (or 2*N*_e_*S*) for diploids (or haploids) represents the strength of selection favoring A/T composition, where negative values of *S* represent selection favoring G/C. S is calculated using the equation, *P*_AT_ = 1/(1 + *m*^−1^*e*^−*S*^), where *P*_AT_ is the fraction of AT nucleotides.

| Species | Type of study | Total number of base-substitution mutations or SNPs | AT mutation bias ($m = v/u$) | Expected equilibrium GC content $[1/(1+m)]$ | Whole-genome GC content | 4-fold site GC content | $S$ (avg) | $S$ (4-fold) |
| --- | --- | --- | --- | --- | --- | --- | --- | --- |
| *P. tetraurelia*<sup>1</sup> | MA | Not known | 0.88 | 0.53 | 0.42 | 0.54 | 0.197 | -0.013 |
| *P. tetraurelia* | population | 37 | 0.95 | 0.51 | 0.42 | 0.54 | 0.163 | -0.047 |
| *P. biaurelia*<sup>2</sup> | MA | 55 | 0.79 | 0.56 | 0.40 | 0.51 | 0.278 | 0.085 |
| *P. sexaurelia*<sup>2</sup> | MA | 87 | 0.49 | 0.67 | 0.39 | 0.47 | 0.504 | 0.368 |
| *P. sexaurelia* | population | 112 | 1.56 | 0.39 | 0.39 | 0.47 | 0.000 | -0.137 |
| *P. caudatum* | population | 96 | 5.02 | 0.17 | 0.22 | 0.07 | -0.151 | 0.397 |
| Other species from previous studies: |  |  |  |  |  |  |  |  |
| *S. cerevisiae*<sup>3</sup> | MA | 13 | 0.00 | 1.00 | 0.17 | 0.08 | >> 0 | >> 0 |
| *Caenorhabditis Elegans*<sup>4,5</sup> | MA | 25 | 3.79 | 0.21 | 0.24 | 0.14 | -0.073 | 0.224 |
| *Daphnia pulex*<sup>6</sup> | MA | 6 | 2.48 | 0.29 | 0.38 | 0.30 | -0.182 | -0.026 |
| *Caenorhabditis Briggsae*<sup>7</sup> | MA | 15 | 18.10 | 0.05 | 0.25 | 0.14 | -0.780 | -0.451 |
| *Drosophila melanogaster*<sup>8</sup> | MA | 28 | 381.25 | 0.00 | 0.18 | 0.06 | -1.923 | -1.386 |

Next, derived singleton alleles at 4-fold degenerate sites were used to quantify the number of G/C to A/T mutations relative to A/T to G/C mutations and thus infer an estimate of AT mutation bias from population data. Again, we find that *m* is close to 1.0 (i.e., no mutation bias towards A/T) in both *P. tetraurelia* and *P. sexaurelia*, with the predicted GC content under mutation equilibrium being remarkably close to that of their 4-fold degenerate sites (Table 1). These results suggest that the composition of 4-fold degenerate sites is mostly determined by mutation. The same can be seen by calculating the strength of selection (*S* = 4*N*_e*S*_) favoring A/T at 4-fold degenerate sites. As *S* usually takes values between 0.1 and 4.0 (Lynch 2007; Long, Sung, et al. 2018) and we obtain much lower magnitudes (0.0-0.5) in the *P. aurelia* species (Table 1), we infer negligible or weak selection at 4-fold degenerate sites, consistent with absence of codon bias at the third position.

In contrast to the *P. aurelia* species, we infer a significant AT mutation bias in *P. caudatum* (Table 1), yielding a predicted GC content at equilibrium of ~17%, with the observed GC content in intergenic and tRNA regions (16% and 20%) being remarkably close to this value. These calculations suggest that genome-wide GC content in *P. caudatum* mitochondria is mostly governed by mutation bias, and that there has been a major shift in mutation bias along the branch leading to the *P. aurelia* species.

### Recombination in *Paramecium* mitochondrial genomes

Next, we sought to evaluate the evidence for the occurrence of recombination in *Paramecium* mitogenomes by performed multiple tests for its presence. First, we evaluated the relationship between linkage disequilibrium (LD), calculated by *r^2^* (Hill and Robertson 1968) and distance between sites; *r^2^* does not decrease with distance in *P. tetraurelia*, *P. sexaurelia*, and *P. caudatum* (*P. multimicronucleatum* isolates lacked sufficient number of polymorphic sites; Supplementary Figure 12), consistent with the expectation under no recombination. Next, we conducted the four-gamete test (FGT) (Hudson and Kaplan 1985), which detects recombination by searching for pairs of polymorphic sites with all four segregating haplotypes and assumes that such pairs of sites must have arisen via recombination (under the infinite-sites model). We found all four gametes at 2 pairs of sites (1.45 × 10^−5^ of all pairs) in *P. tetraurelia*, 312781 pairs (2.14 × 10^−2^ of all pairs) in *P. sexaurelia*, 42647 pairs (1.21 × 10^−2^ of all pairs) in *P. caudatum*, and 0 pairs of sites in *P. multimicronucleatum*. The observed variation between species correlates well with levels of total sequence diversity and indicates the bias in power to detect recombination towards species with more sequence variation. Although results from the FGT suggest the presence of some recombination, the probability of finding all four gametes does not increase with the distance between the pairs of sites (slope= −7.64 × 10^−9^, *p* = 0.765 for *P. sexaurelia*; slope = −1.39 × 10^−8^, *p* = 0.792 for *P. caudatum*), contrary to the expectation under recombination, and is inconsistent with other distance-based analyses. Violation of the infinite-sites model (i.e., recurrent mutation at the same site) can result in the presence of pairs of sites with all four alleles. The FGT works best for species and genomes whose recombination rate is much larger than the mutation rate, which may not be the case in *Paramecium* mitochondria.

In order to test if the presence of four gametes was more likely to be caused by recombination or mutation, we used LDhat (McVean, et al. 2002) to test for recombination and estimate 2*N_e_r*, using values of 2*N_e_μ* estimated by nucleotide diversity as calculated in Johri et al., 2017 (Supplementary Table 4). Here *r* is the recombination rate, *μ* is the mutation rate and *N_e_* is the effective population size of the species under consideration. 2*N_e_r* estimated under the finite-sites mutation model (Wakeley 1997; Hudson 2001) was found to be 0.0 for all four species, and is consistently 0.0 in all non-overlapping windows (1000 bp) spanning the genome. We also used permutation tests (McVean, et al. 2002) to detect recombination. The idea behind these tests is that in the absence of recombination the relative position of SNPs would not change inferred values of statistics used to measure recombination. Thus, the absence of recombination, statistics like sum-of-distance between sites with all four gametes (G4) (Meunier and Eyre-Walker 2001), and correlation between *r^2^* or *D*′ and physical distance, would not be significantly different from an expectation obtained by randomly shuffling SNPs and computing these statistics. We find no statistical significance for the presence of recombination using permutation tests (Supplementary Table 4) and thus conclude that there is no recombination in *Paramecium* mitochondria.

### Efficacy of selection in mitochondria versus the nucleus

Finally, we asked if mitochondrial genes experience weaker efficacy of purifying selection compared to nuclear genes, as would be expected due to smaller effective population size and the lack of recombination in the mitochondria. We did so by comparing multiple statistics in each of the two genomes. Because our statistics included divergence at synonymous sites, we conducted these analyses primarily on *P. tetraurelia* where values of d*S* were calculated with respect to the closest outgroup species *P. biaurelia*, for which we had available sequences in both the nucleus and mitochondria. Average d*S* values for the set of nuclear and mitochondrial genes were found to be 0.885 and 1.174 respectively (Table 2), a small but significant difference. In order to control for differences in d*S* driving the patterns, we also conducted all analyses restricted to genes with d*S* < 1.0. For the set of genes with d*S* < 1.0, the average value of dS is not significantly different between the two genomes, 0.636 among the nuclear and 0.658 among the mitochondrial genes. A potential caveat of comparing the efficacy of purifying selection between all genes present in the mitochondria vs. nucleus, is that we might instead be measuring differences in strength of purifying selection. In order to correct for that, we also compared nuclear genes with similar functions to those in the mitochondria. We compared the 14 mitochondrial genes with ~13-27 nuclear genes involved in the oxidative phosphorylation pathway, as well as the 8 ribosomal genes in the mitochondria with ~77-430 nuclear genes that encode for structural components of the ribosomes.

**Table 2:**
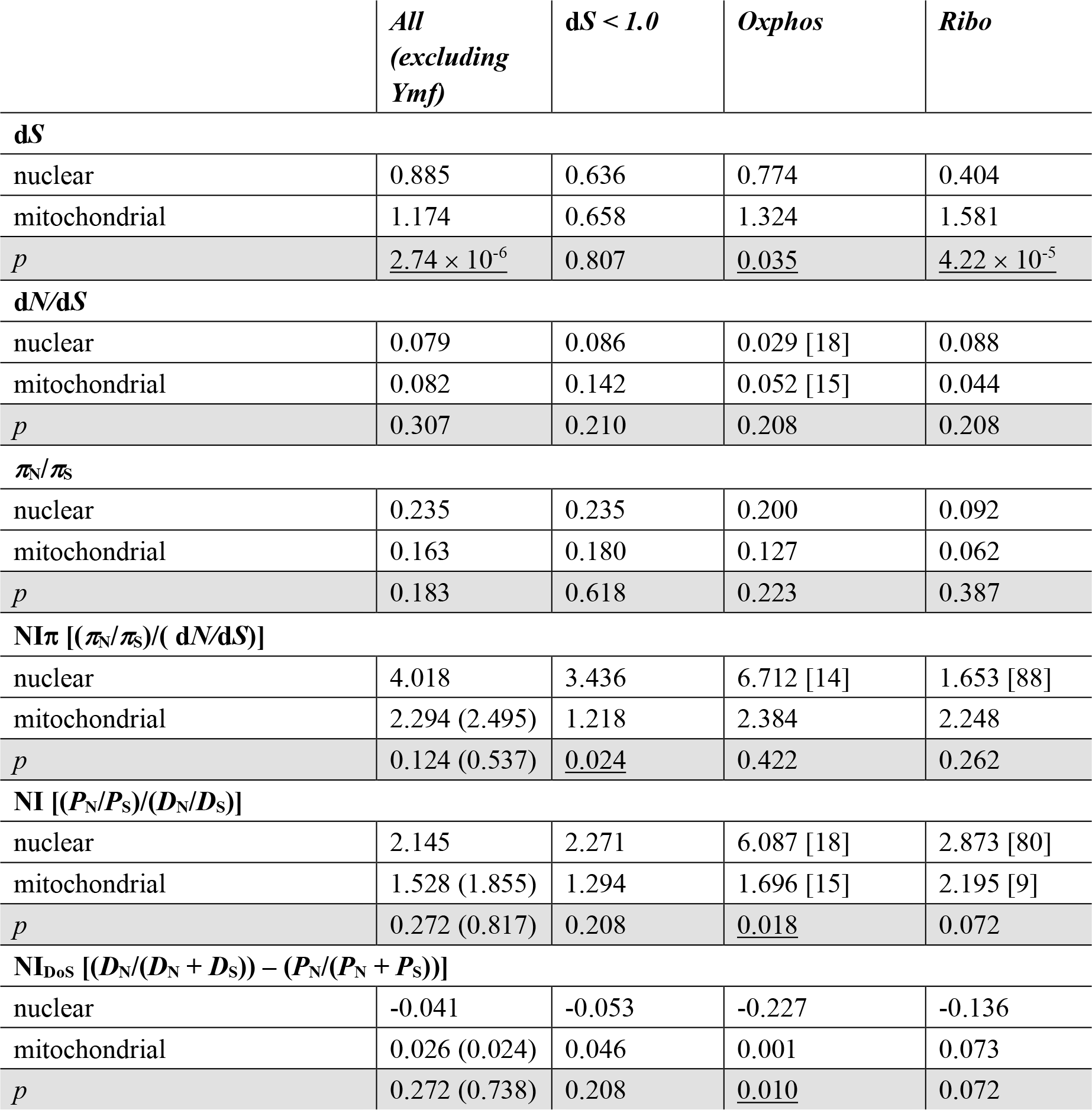
Neutrality index (NI) of mitochondrial and nuclear genes in *P. tetraurelia*, calculated using multiple statistics. All *p* values (corrected for multiple testing by Holm-Bonferroni method) represent comparisons between nuclear and mitochondrial statistics. Statistically significant scores (*p* < 0.05) are underlined.

Average d*N*/d*S* is similar or slightly lower for genes in the nucleus (mt = 0.082, nuc = 0.079, *p* = 0.307; Table 2), with the exception of the well-conserved ribosomal genes, which have lower d*N*/d*S* in the mitochondria (mt = 0.044, nuc = 0.088, *p* = 0.208). None of these differences in d*N*/d*S* are significant, suggesting that the average amount of purifying selection experienced by genes in the nucleus is not significantly different from that experienced by those in the mitochondria.

The efficacy of purifying selection can also be measured by estimating the fraction of segregating nonsynonymous polymorphisms that undergo fixation. Such a measure can be calculated using the neutrality index (NI = (*P*_*n*_ / *P_s_*) / (*D*_*N*_ / *D*_*S*_), where *P*_*n*_ and *P_s_* are the number of nonsynonymous and synonymous polymorphisms respectively; *D*_*N*_ and *D*_*S*_ are the number of nonsynonymous and synonymous fixed differences respectively). The neutrality index is usually found to be larger than 1.0 for genes experiencing purifying selection because mildly deleterious variants are allowed to segregate among individuals, but rarely fix in populations. Higher values of the NI suggest that smaller proportion of segregating nonsynonymous variants are allowed to fix in the population, indicating stronger efficacy of purifying selection. The number of branch-specific substitutions at synonymous (*D*_*S*_) and nonsynonymous (*D*_*N*_) sites was calculated by ancestral reconstruction, using *P. tetraurelia*, *P. biaurelia*, *P. sexaurelia*, *P. caudatum*, and *P. multimicronucleatum*, for both the nucleus and the mitochondria, and restricted to sites for which changes could be confidently inferred along specific branches. The advantage of using ancestral reconstruction to infer *D*_n_ and *D*_s_ is that particular sites that are very fast evolving can be excluded from the analysis, thus minimizing problems arising due to saturation of divergence at synonymous sites. Comparing the distributions of values of NI in the mitochondrion and nucleus shows that overall there is no significant difference between the two sets when all genes are included (whether we include or exclude *Ymf* genes) in *P. tetraurelia* (Table 2). However, for genes involved in oxidative phosphorylation, nuclear genes appear to be experiencing significantly stronger efficacy of purifying selection than mitochondrial genes (mt NI = 1.696, nuc NI = 6.087, *p* = 0.02).

We also calculated the neutrality index as (*π*_n_/*π*_s_)/ (*dN/dS*) (Betancourt, et al. 2012; denoted by NIπ below) for each gene with *dN/dS* obtained from pairwise comparison with respect to *P. biaurelia*. In this case all sites contribute to the analysis, but the maximum likelihood estimate of pairwise d*N*/d*S* can minimize biases due to saturation of d*S*. Again, we observe that neutrality index of nuclear genes is either similar or higher than those of mitochondrial genes. Genes involved in oxidative phosphorylation in the nucleus have higher but not significantly different NIπ compared to those in the mitochondria (mt = 2.384, nuc = 6.712, *p* = 0.422).

In order to minimize statistical biases that can arise due to sampling, we also estimated a variation of the neutrality index NI_DoS_ = *D*_n_/(*D*_n_+*D*_s_) - *P*_n_/(*P*_n_+*P*_s_) (Stoletzki and Eyre-Walker 2011), where more negative values represent stronger efficacy of purifying selection. As previously observed, we find no significant difference between the neutrality index of genes in the nucleus vs. mitochondria except for Oxphos genes, where nuclear genes have significantly lower values of NI_DoS_ than mitochondrial genes (mt NI_DoS_ = 0.001, nuc NI_DoS_ = −0.227, *p* = 0.01) This suggests that there are more deleterious variants segregating in the nucleus than in the mitochondrion in *P. tetraurelia*. Overall, mitochondria appear to experience either similar or stronger efficacy of purifying selection than the nucleus.

Lastly, we used *π*_n_/*π*_s_ as a proxy for the efficacy of recent purifying selection. This comparison can be extended to all four species (Supplementary Table 5). We find that average *π*_n_/*π*_s_ of mitochondrial (mt) and nuclear genes (nuc) is not significantly different in *P. tetraurelia* (mean mt = 0.163; mean nuc = 0.235; *p* = 0.18) and *P. caudatum* (mean mt = 0.135; mean nuc = 0.170; *p* = 0.22) respectively. However, mean *π*_n_/*π*_s_ in the mitochondrial genes is significantly lower than that of nuclear genes in *P. sexaurelia* (mean mt = 0.051; mean nuc = 0.268; *p* < 2.2 × 10^−16^) and *P. multimicronucleatum* (mean mt = 0.099; mean nuc = 0.178; *p* = 1.87 × 10^−3^) respectively, suggesting that mitochondrial genes might be under stronger purifying selection than nuclear genes.

As side note, values of NI for *P. sexaurelia*, *P. caudatum* and *P. multimicronucleatum* were consistently found to be much less than 1.0, likely due to under-estimation of changes at synonymous sites. We aimed to reduce the bias caused by saturation of *D*s by re-calculating NI only in the mitochondrion using all 13 taxa (in order to break up longer branches). We continue to recover extremely low values of NIs (Supplementary Table 6) and fail to obtain values close to those obtained using NI*π*, suggesting that the absolute values of NI can be misleading and have to be interpreted with caution.

## DISCUSSION

In this study, we greatly expand the set of sequenced ciliate mitochondrial genomes by presenting a wider sampling of the mitogenome diversity within the *Paramecium* genus. Using this wealth of sequence data, we characterize the diversity and conservation of genome organization and gene content, and we study in depth the population genetic characteristics such as mutational and selection pressures acting on mtDNA within the genus.

### Ciliate-specific mitochondrial genes

*Paramecium* mitochondrial genomes possess 16 lineage-specific open-reading frames (referred to as *Ymf* genes) that have no known homologs in non-ciliate species and lack assigned functions, but are nonetheless conserved across *Paramecium* and *Tetrahymena* species. Other ciliates like *Oxytricha* have also been found to have unidentified open-reading frames (Swart, et al. 2012), but not all *Ymf* genes have homologs identified in ciliates outside of *Oligohymenophora* (which contains both *Tetrahymena* and *Paramecium*). We find that *Ymf* genes are evolving faster, mostly due to relaxed purifying selection, as Ymf genes are not significantly more likely to undergo positive selection than other genes. Shedding light on the identity of the *Ymf* genes could possibly indicate entirely new sets of genes present in the ancestor of *Oligohymenophorean* mitochondrial genome.

We found that parts of five of the 16 *Ymf* genes show homology to standard mitochondrial proteins, especially ribosomal proteins. As ciliates are evolutionarily distant from most model organisms in the eukaryotic tree, it is possible that some *Ymf* genes are ribosomal subunits or genes belonging to the NADH complex and are simply not identifiable because of extreme divergence. Similar findings have been reported for other protozoan mitochondrial genomes in the past (e.g. de Graaf, et al. 2009; Pombert, et al. 2013; Burger, et al. 2016; Skippington, et al. 2017) that have diverged substantially from the most well-studied mammalian mitochondrial genomes. In some species, RNA editing (including insertions and deletions) can be substantial (Gray 2003), which can mask the proteins encoded from identification from genomic sequence data. We however found no evidence of RNA editing, consistent with a previous report in *P. tetraurelia* mitochondria (Orr, et al. 1997), thus it is unlikely to account for the observed divergence. An interesting possibility is that most of the *Ymf* genes are either part of or interact with the ATP synthase complex. In *Tetrahymena thermophila*, the ATP synthase has been reported to form an unusual structure possessing completely novel subunits whose orthologs are not identifiable in other organisms, some of which are *Ymf* genes (Balabaskaran Nina, et al. 2010). It is therefore possible that ciliates possess structurally unique proteins that perform relatively conventional functions in the mitochondria, but are difficult to identify based on other known sequences. Although the origin of these genes is unclear, it is intriguing that a set of such fast evolving genes is preserved across highly diverged species.

### Change in mutation spectrum and nucleotide composition

We used a combination of previous mutation-accumulation studies and our population-genomic data to determine that the change in nucleotide composition of *P. aurelia* mitochondrial genomes toward higher GC is most likely the result of changes in mutational biases. Unfortunately, due to our modest sample sizes, some of the SNPs observed as singletons in our data may in fact be fairly common in the population at large. Our estimates of mutation spectra might thus be biased by selection. Mutation-accumulation lines in *P. caudatum* would help further disentangle these forces. Because ciliates have very low mutation rates, their mutation-accumulation study requires a large number of generations to produce only a handful of mutations, making it very difficult to obtain precise estimates of the mutation spectrum. The most feasible strategy to more precisely estimate these spectra in *Paramecium* would thus be to acquire larger population samples in order to observe lower-frequency variants. We also note that although an alternative explanation for higher GC content in the *P. aurelia* species could be biased gene conversion, we found no evidence of mitochondrial recombination (including non-crossovers) in the *P. aurelia* species.

An interesting question raised by our results is how fast the mutation spectrum in mitochondria evolves across species (Lynch, et al. 2008; Montooth and Rand 2008). Re-analyzing data from previous work on mutation-accumulation lines in mitochondria of other model organisms presents a number of interesting observations (Table 1). First, within opisthokonts, there is a huge variation in mutation bias (*m*) in the mitochondria, ranging from nearly 0 (strongly biased towards GC) to values much larger than 1 (biased towards AT; Table 1). However, most species except *S. cerevisiae* have a mutation bias towards A/T, consistent with most mitochondrial genomes being AT-rich. Second, across opisthokonts, we find that selection can have a significant impact on mitochondrial GC content. The effect of selection (*S*) can be observed by comparing observed genome-wide nucleotide composition (or specifically at 4-fold degenerate sites) with the expected composition under mutation equilibrium (Table 1). Again, in most species selection favors higher G/C, but in *S. cerevisiae* there appears to be strong selection favoring A/T genome-wide. Thus, there might not be a universal direction for mutation bias or selection for nucleotide composition in mitochondria. Finally, closely related species in other lineages have been shown to have significant differences in mutation bias in the mitochondria as observed for example between *C. elegans* and *C. briggsae* (mutation bias re-calculated by combining observations from Howe, et al. 2010; Konrad, et al. 2017). All of these observations suggest that it may not be that such an unusual event for the mutational biases to shift away from AT along the long branch leading to the *P. aurelia* species. However, the exact nature of the biochemical mechanisms responsible for remains an intriguing open question for future research.

### Similar efficacy of purifying selection experienced by the mitochondria and nucleus

Mitochondrial genomes are often non-recombining and are usually passed on via uniparental inheritance. Mitochondria are therefore expected to have lower effective population sizes than that of the nucleus within the same organism (Lynch and Blanchard 1998; Neiman and Taylor 2009). One consequence of reduced effective population sizes is an increased probability of segregation and fixation of slightly deleterious mutations. There have been multiple contradictory reports on whether mitochondrial genes experience stronger or weaker purifying selection than the nuclear genes. Studies examining a small number of protein-coding loci across a large number of species have concluded that mitochondria experience less effective purifying selection than the nucleus (Weinreich and Rand 2000; Betancourt, et al. 2012; Popadin, et al. 2013). However a recent study conducted on multiple individuals and whole genomes in *Drosophila melanogaster* and humans found no significant difference between the efficacy of purifying selection in the mitochondrial vs. nuclear protein-coding genes (Cooper, et al. 2015).

We confirm the absence of recombination in mitochondria of three species of *Paramecium*. Previous studies in *P. tetraurelia* (Adoutte, et al. 1979; Barth, et al. 2008) and *P. primaurelia* (Beale, et al. 1972) had reached similar conclusions using a small number of markers. We also evaluated the extent of purifying selection in ciliate mitochondria relative to the nucleus primarily for *P. tetraurelia*. We find that despite the lack of recombination in the mitochondria, the efficacy of selection in the mitochondria is similar to if not stronger than that of the nucleus.

Our results seem to be in discordance with theoretical predictions according to which one would expect the efficacy of selection in the mitochondria to be lower than in the nucleus. One possibility is that we observe stronger efficacy of selection in the mitochondria because the magnitude of negative selection itself is stronger in the mitochondrial genes (Popadin, et al. 2013), which differ from the average nuclear gene in several key ways (Adrion, et al. 2016). Most mitochondrial genes are expressed at higher levels compared to nuclear genes (Havird and Sloan 2016). Mitochondrial encoded proteins (*cox1* and *cox2*) that are part of the oxidative phosphorylation (OXPHOS) pathway are core enzyme catalytic subunits (Tsukihara, et al. 1996; Zhang and Broughton 2013; Havird and Sloan 2016). Finally, most genes retained in the mitochondria encode for highly hydrophobic proteins and have a high GC content (Johnston and Williams 2016). Thus, finding comparable sets of genes between these two genomes is admittedly difficult (but see Lynch 1996; Lynch 1997).

Lastly, our results need to be evaluated in the light of *Paramecium*-specific life cycle. In *Paramecium*, there is almost no exchange of cytoplasm during conjugation (Koizumi and Kobayashi 1989; Meyer and Garnier 2002). Thus it appears that both parents pass on their mitochondria to their respective offspring, without exchange or degradation of mitochondrial genomes. In addition, mitochondria double before binary fission and are distributed randomly across the daughter cells. Thus, mitochondria do not appear to undergo bottlenecks during any stage of the life cycle in *Paramecium*, and experience no associated reduction in effective population size. On the other hand, *Paramecium* species frequently undergo asexual reproduction, thereby reducing the nuclear effective population size. Therefore, effective population sizes of mitochondrial and nuclear genomes may be more similar in *Paramecium* than in many other organisms. Due to the combination of sexual and asexual reproduction, under equilibrium conditions, expected heterozygosity in the nucleus would take values between 4*N*_e_*μ*_n_ - 2*N*_e_*μ*_n_. Similarly, in the mitochondria, expected hetrozygosity would be ~2*N*_e_*μ*_m_. Assuming that the ratio of divergence at silent sites can be used as a proxy for the ratio of mutation rate between the two compartments, it is possible to approximately estimate the ratio of effective population sizes of the two compartments as *N*_e(m)_/*N*_e(n)_ = *y* × (*π*_m_/*π*_n_)/(*d*_m_/*d*_n_), where *y* would be some number between 1 and 2. For *P. tetraurelia*, for which we have relatively closer outgroup species and thus more reliable divergence estimates, our estimated range of *N*_e(m)_/*N*_e(n)_ is 0.94 − 1.88. Although underestimation of neutral divergence in the mitochondria relative to the nucleus could skew our inference slightly, the above calculation suggests that effective population sizes of mitochondria may be similar or, larger than that of the nucleus in *Paramecium*.

We therefore conclude that our finding of similar or stronger efficacy of selection in the mitochondria relative to the nucleus in *Paramecium* may lie within theoretical expectations given *Paramecium*’s unique life cycle and mode of mitochondrial transmission. A better understanding of the *Paramecium* life cycle in the wild might help build more appropriate null expectations in the future. Our results suggest the possibility that unicellular eukaryotes in general may have larger mitochondrial than nuclear effective population sizes and more efficacious purifying selection in the mitochondria might be more common than believed.

## METHODS

### Genome sequencing and assembly

Single isolates of *P. jenningsi* (strain: M), *P. octaurelia*, *P. decaurelia* (strain: 223), *P. dodecaurelia* (strain: 274), *P. novaurelia* (strain: TE) and *P. quadecaurelia* (strain: 328) were used to extract macronuclear DNA. DNA extraction, sequencing library preparation and genome sequencing were previously described (Johri et al. 2017). Sequencing reads were assembled using SPAdes (Bankevich, et al. 2012; version 3.5.0) after removing potential adapter sequence with Trimmomatic (Bolger, et al. 2014; version 0.33). Mitochondrial contigs were identified from the resulting assemblies by BLAST (Altschul, et al. 1997) searches against the published *P. caudatum* and *P. tetraurelia* mitochondrial genomes. Mitochondrial re-sequenced data from 5-10 isolates of *P. tetraurelia*, *P. sexaurelia*, *P. caudatum* and *P. multimicronucleatum* were downloaded from SRA (PRJNA490059) and SNPs were called as described in Johri et al. 2017, using reference genomes of strain 99 for *P. tetraurelia*, strain 130 for *P. sexaurelia*, C104 for *P. caudatum*, and M04 for *P. multimicronucleatum*.

### Mitochondrial genome annotation

Genome annotation was carried out as follows. Protein coding genes were identified by generating all ORFs longer than 60aa in all six frames, using the Mold, Protozoan, and Coelenterate Mitochondrial Code (i.e., UGA codes for W instead of being a stop codon) and all alternative start codons specific to *Paramecium* (AUU, AUA, AUG, AUC, GUG, and GUA), and retaining the longest ORFs associated with each stop codon. BLASTP was then used to identify homologs of annotated mitochondrial proteins in *P. tetraurelia* and *P. caudatum*. Additional ORFs were identified by imposing the requirement that their length exceeds 100aa, and subsequently annotated using BLASTP against the Non-redundant protein sequences (nr) database and HMMER3.0 (Eddy 2011) scans against the PFAM 27.0 database (Finn, et al. 2014). tRNA genes were annotated with tRNAscan-SE (Schattner, et al. 2005; version 1.21), using the “Mito/Chloroplast” source. rRNA genes were identified using Infernal (Nawrocki, et al. 2009; version 1.1.1).

### Identification of telomeres

Telomeric repeats were identified as follows. The first and the last 200bp of each *de novo* assembled mitochondrial genome were used as input to the MEME *de novo* motif finding program (Bailey, et al. 2009; version 4.6.1), which was run with the following parameters: “- *maxw 25 -dna -nmotifs 5 -mod* anr”. The repetitive units defined that way were then manually aligned to each other and refined to arrive at final telomeric repeats comparable across all species.

### RNA-seq analysis

For each species, sequencing reads were aligned against a combined Bowtie (Langmead, et al. 2009) index containing both the nuclear and mitochondrial genomes using TopHat2 (Kim, et al. 2013; version 2.0.8) with the following settings: *“--bowtie1 --no-discordant --no-mixed -- microexon-search --read-realign-edit-dist 0 --read-edit-dist 4 --read-mismatches 4 --min-intron-length 10 --max-intron-length 1000000 --min-segment-intron 10 --min-coverage-intron 10*”. Custom python scripts were then used to identify sequence variants relative to the mitogenomes assemblies.

### Building phylogenetic trees

Nucleotide sequences were extracted, aligned using MUSCLE, and concatenated. Missing data was encoded as “N”. RAxML was used to build the tree with GTRGAMMA as the substitution model. Boostrap values for 1000 replicates were obtained via the fast method recommended by RAxML with the following command line:

*raxmlHPC -f a -s sequences.fasta −n sequences_boot -m GTRGAMMA -T 50 -p 31 -x 7777 -N 1000*

### Estimation of *dN/dS*, *D*_n_, *D*_s_, *P*_n_, *P*_s_, and *π*_n_/*π*_s_

*dN*/*dS* was estimated either across the phylogeny using CODEML, PAML (Yang 2007; version 4.9a) and for the *P. aurelia* species it was also estimated pairwise w.r.t the closest outgroup species using yn00, PAML. *π*_n_/*π*_s_ was obtained for all protein coding genes in the 4 species-*P. tetraurelia*, *P. sexaurelia*, *P. caudatum*, and *P. multimicronucleatum*, where total number of changes in synonymous sites was greater than 1. This filter was executed in order to reduce errors in *π*_n_/*π*_s_ due to very low values of synonymous polymorphisms in a gene.

*D*_n_ and *D*_s_, the number of nonsynonymous and synonymous changes, were inferred by performing ancestral reconstruction at each site, and then counting branching-specific substitutions. Ancestral reconstruction (PAML, baseml, GTR model) was conducted over the phylogeny of all available taxa. For comparing statistics between mitochondrial and nuclear genes, the ancestral reconstruction was performed over the same set of taxa for both, i.e. over *P. tetraurelia*, *P. sexaurelia*, *P. caudatum*, and *P. multimicronucleatum*. For all analyses that involved ancestral reconstruction, only sites whose posterior probability of the inferred ancestral state ≥ 0.85 were used in the analyses—this filter was imposed for counting synonymous and nonsynonymous polymorphisms (*P*_*s*_ and *P*_*n*_, respectively) as well as divergent sites (*D*_*n*_ and *D*_*s*_). There may be some concern that ancestral reconstruction could end up biasing the ratio of *D*_n_/*D*_s_ as many more changes at synonymous sites might result in lower confidence in inferring ancestral states at synonymous but not nonsynonymous sites. Such a bias would increase *D*_n_ relative to *D*_s_ and thus decrease values of NI. Therefore, all analyses involving *D*_n_ and *D*_s_ were also performed including all sites, with no filter, and results remained unchanged.

### Calculation of multiple estimators of Neutrality Index (NI) and statistical tests

Several estimators of neutrality index have been proposed in order to counter different biases. The estimators we used were calculated as follows:

Simple neutrality index, NI = *P*_n_/*P*_s_/*D*_n_/*D*_s_ (Rand and Kann 1996).
NI_π_ = *π*_n_/*π*_s_/*d*_N_/*d*_S_, where *d*_N_/*d*_S_ was calculated pairwise, w.r.t closest outgroup species (Betancourt, et al. 2012).
NI_TG_ = ∑_i_[*D*_si_.*P*_ni_/(*P*_si_ + *D*_si_)] / ∑_i_ [*P*_si_.*D*_ni_/(*P*_si_ + *D*_si_)], where *i* is the *i*th gene (Tarone 1981; Greenland 1982).
NI_DoS_ = *D*_n_/(*D*_n_+*D*_s_) - *P*_n_/(*P*_n_+*P*_s_) (Stoletzki and Eyre-Walker 2011).

For McDonald Kreitman test (McDonald and Kreitman 1991), Fisher’s exact test in R (R-Core-Team 2014) was used to test significance. In all cases p value was corrected by Bonferonni-Holm method (Holm 1979) for multiple tests, using R.

### Mutation spectrum from mutation accumulation (MA) lines

MA line experiments for *P. tetraurelia* had previously been published (Sung et al. 2012) and results from their analysis were used directly in this study.

MA experiments carried out in order to obtain nuclear mutation rates in *P. biaurelia* and *P. sexaurelia* (Long, Doak, et al. 2018) were reanalyzed as follows. Sequencing reads were assembled for each MA line individually and mitochondrial contigs identified as described above. A composite consensus mitochondrial genome sequence was then created from the individual assemblies by creating multiple sequence alignments of all mitochondrial contigs using MAFFT (Katoh and Standley 2013; version 7.221) and retaining the most frequent base for each alignment column (with the exception of telomeres, which were manually curated). Adapter-trimmed reads were then aligned in a 2×100bp format against a combined Bowtie index, containing a combination of the nuclear and consensus mitochondrial genomes, allowing for up to 3 mismatches and retaining only unique reads. Putative mutations were identified by requiring that any variant is supported by at least 3 non-redundant read pairs on each strand, is supported by not more than 4 times more reads on one strand than on the other, and is also observed in ≥5% of reads covering a given position. Telomeric sequences were excluded due to an excessively high number of sequence variants observed in those regions.

### Mutation spectrum from population genomics data

For each SNP, the ancestral allele was inferred by performing ancestral reconstruction on the 13-taxa phylogeny to predict the nucleotides on internal nodes (see above). The ancestral allele was used to infer the derived allele segregating in *P. tetraurelia*, *P. sexaurelia*, *P. caudatum* and *P. multimicronucleatum*. This analysis was restricted to sites where the ancestral nucleotide was inferred with confidence score >= 0.90, where the derived allele was present in only a single individual, i.e. was a singleton, and was at 4-fold degenerate site. Of these, we counted all mutations that were from G/C to A/T or from A/T to G/C, with respect to the total number of utilizable sites that were counted according to the same criteria as above. Care was also taken to remove all sites that were part of overlapping ORFs.

### Calculation of bias in mutation spectrum

Mutation rate of A/T ⟶ G/C (*u*) and G/C ⟶ A/T (*v*) mutations was calculated as follows:

*u* = (number of A/T ⟶ G/C mutations) / (total number of utilizable A/T sites)
*v* = (number of G/C ⟶ A/T mutations) / (total number of utilizable G/C sites)

Mutation bias towards A/T (*m*) was calculated as, *m* = *v/u* and the expected equilibrium G/C content was calculated as 1/(1+*m*) following Lynch 2007.

*S* = 4*N*_e*S*_ (or 2*N*_e*S*_), is the population-scaled strength of selection towards A/T nucleotides and can be calculated using the equation, *P*_AT_ = 1/(1 + *m*^−1^*e*^−*S*^), where *P*_AT_ is the observed fraction of A/T sites and *s* is the average selective advantage of A/T over G/C nucleotides (Bulmer 1991). Thus *S* = -ln[(*m*.(1-*P*_AT_))/*P*_AT_] = ln(*P*_AT_/*P*_GC_/*v*/*u*).

### Recombination analyses

All analyses to detect recombination were restricted to SNPs that were biallelic, homozygous, and had a known ancestral state. The statistic (*r*^2^) to measure linkage disequilibrium was calculated as *r*^2^ = (*f*_Aa_ − *f*_A_×*f*_a_)^2^ / [*f*_A_×*f*_a_×(1−*f*_A_)×(1−*f*_a_)]. The program “pairwise” in LDhat 2.2 (McVean, et al. 2002) was used to infer recombination rates using the permuted composite likelihood test as well other permutation tests. These tests were performed under both the gene conversion (average tract length: 500) and cross-over models with 2 values of *θ* (= 4*N*_e_*μ*) for each species: the closest allowed *θ* value lower than that estimated from nucleotide diversity values, and the closest higher value.

### Identifying nuclear genes belonging to Oxidative Phosphorylation pathway and Ribosomal complex

KEGG (Kanehisa, et al. 2017) was used to obtain the full list of genes that are part of complexes involved in Oxidative Phosphorylation (Complexes I-V) for *P. tetraurelia*. A total number of 87 genes in *P. tetraurelia* were obtained, and their corresponding orthologs were identified in other species. For genes encoding proteins that are part of the ribosomal complex, we used the PANTHER (Mi, et al. 2017) annotation obtained in a previous study (McGrath, Gout, Johri, et al. 2014) for all species and selected all genes that were structural constituents of ribosome. This allowed us to start with a set of 585 genes in *P. tetraurelia*, 564 in *P. sexaurelia*, and 213 genes in *P. caudatum*. However, analyses requiring orthologs from all 3 species were conducted with a smaller subset of genes.

## ACKNOWLEDGEMENTS

We thank Jean-Francois Gout for technical help, as well as for a critical reading of the manuscript. We thank Hongan Long for providing us with sequencing data for *P. biaurelia* and *P. sexaurelia* nuclear mutation-accumulation experiments. We also thank Jeffrey Palmer for critical reading of the manuscript and discussions about the project. This work was financially supported by the National Science Foundation (MCB-1050161 and DEB-1257806).

## DATA AVAILABILITY

Assembled mitogenomes and protein coding gene annotations can be accessed at https://github.com/georgimarinov/Paramecium_mitochondrial_genomes.

(Xu, et al. 2012), (Haag-Liautard, et al. 2008),

## SUPPLEMENTARY MATERIALS

**Supplementary Figure 1:**
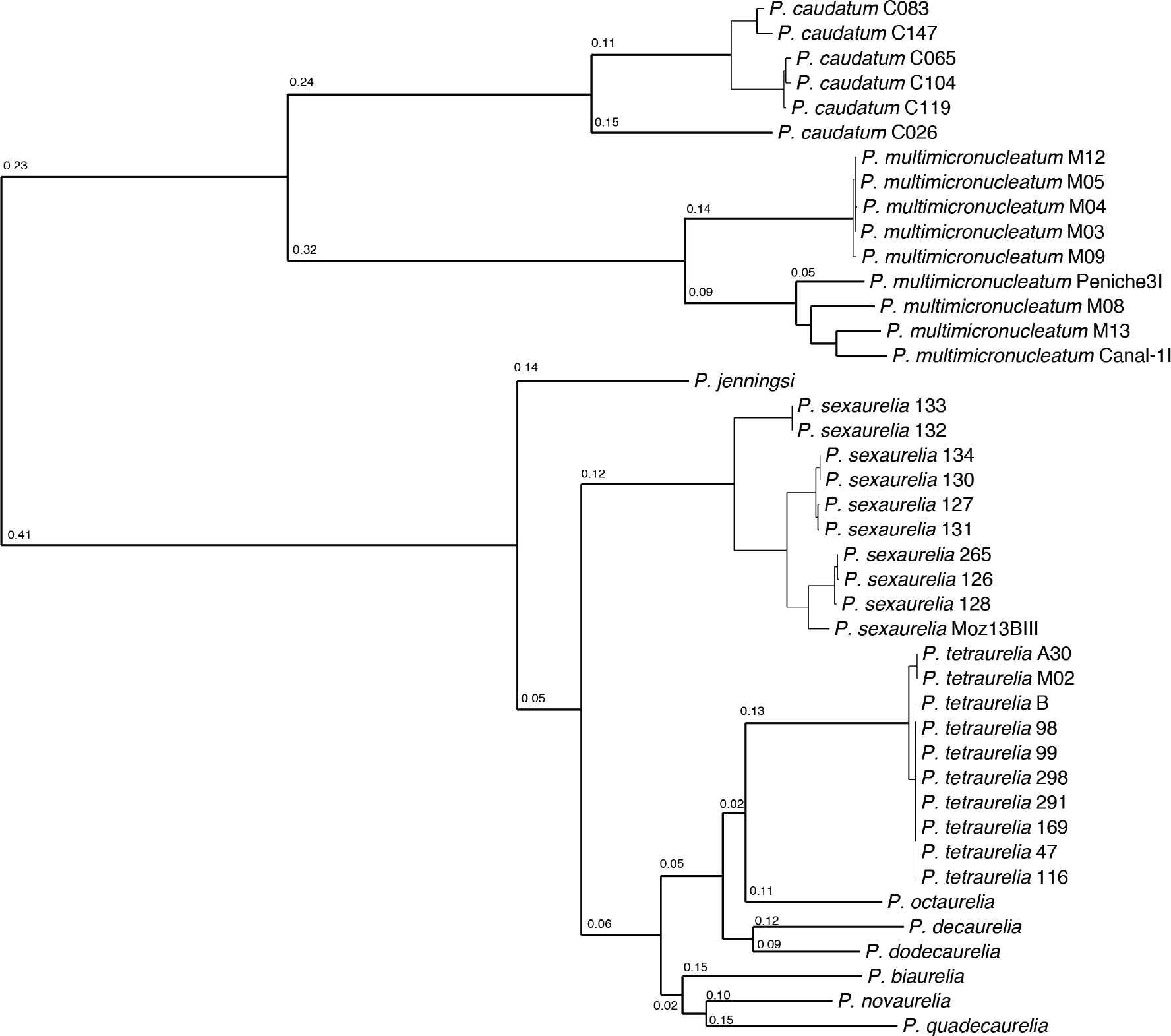
Mitochondrial phylogeny of the *Paramecium* genus including individual isolates within each species. The phylogeny was built using RAxML (version 8.2.11), under the substitution model GTRGAMMA. Numbers on branches indicate total number of substitutions per site.

**Supplementary Figure 2:**
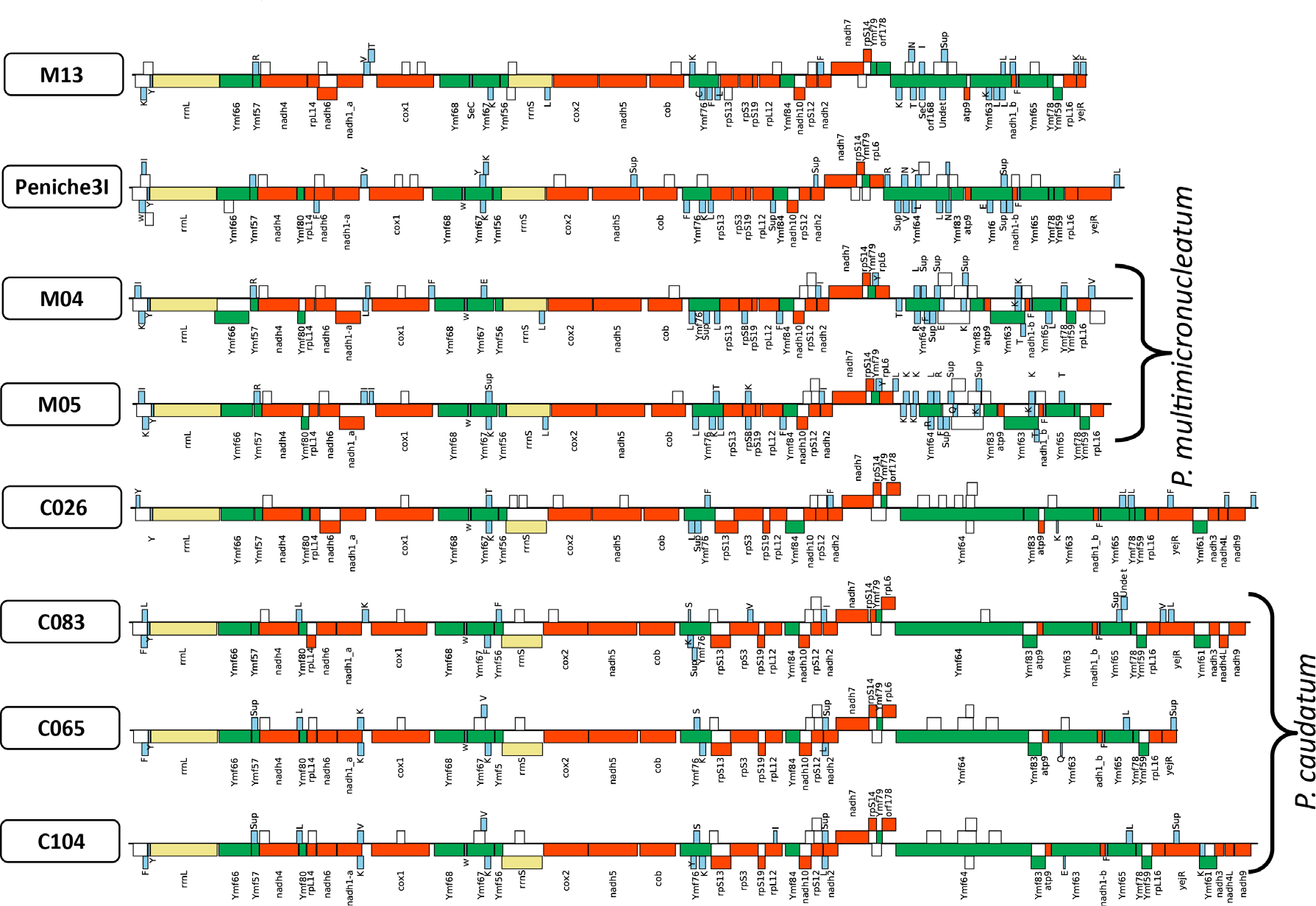
Mitochondrial genomes of isolates belonging to the *P. caudatum* and *P. multimicronucleatum* outgroup lineages. Red: protein coding genes; white: novel predicted ORFs longer than 100aa; yellow: ribosomal RNAs; blue: tRNAs. Top layers: forward-strand genes; bottom layers: reverse-strand genes.

**Supplementary Figure 3:**
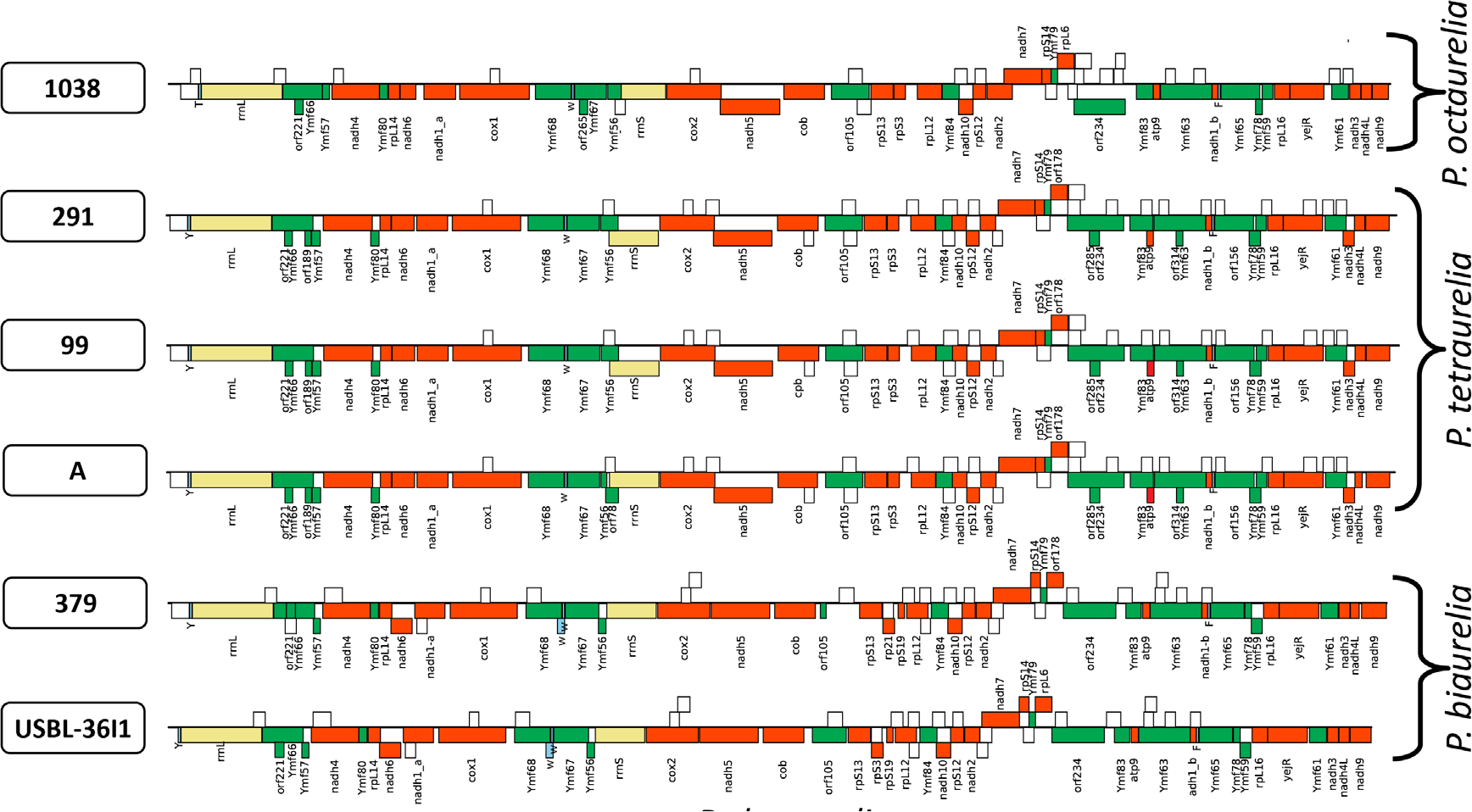
Mitochondrial genomes of isolates belonging to the *P. aurelia* species complexes (except for *P. sexaurelia*, shown in the next figure). Red: protein coding genes; white: novel predicted ORFs longer than 100aa; yellow: ribosomal RNAs; blue: tRNAs. Top layers: forward-strand genes; bottom layers: reverse-strand genes.

**Supplementary Figure 4:**
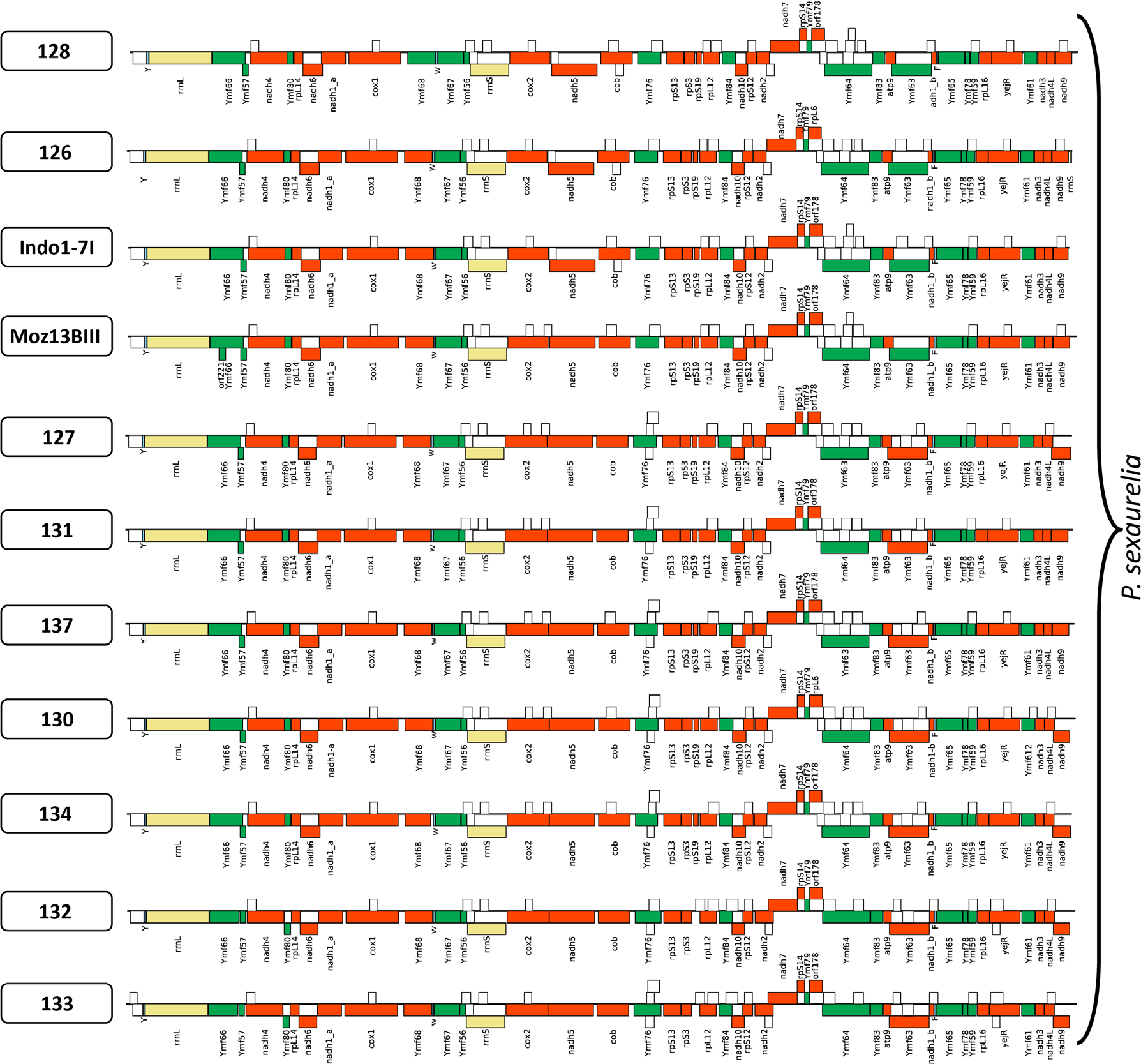
Mitochondrial genomes of isolates belonging to *P. sexaurelia*. Red: protein coding genes; white: novel predicted ORFs longer than 100aa; yellow: ribosomal RNAs; blue: tRNAs. Top layers: forward-strand genes; bottom layers: reverse-strand genes.

**Supplementary Figure 5:**
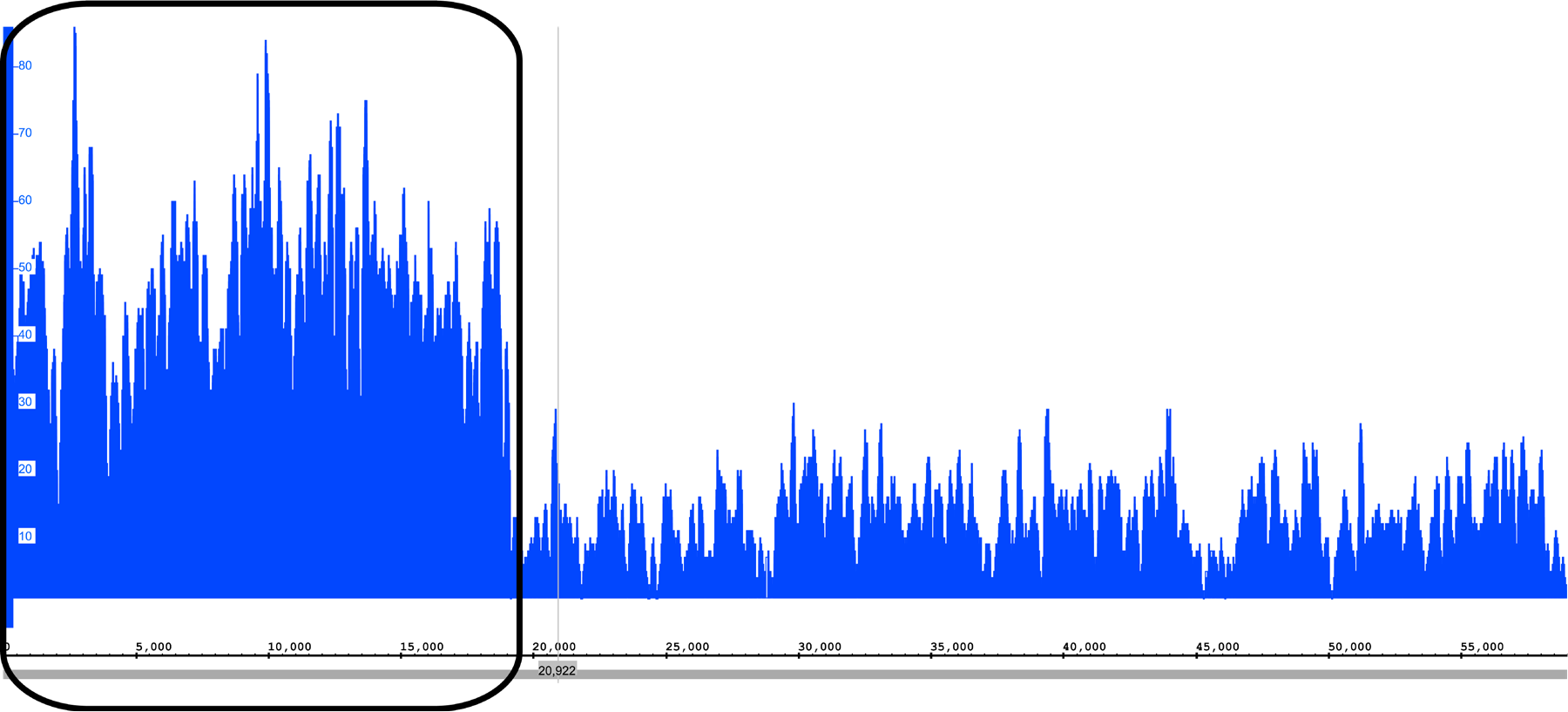
Raw mitogenome assembly and sequencing read coverage for *P. novaurelia*. An ~19kb 3′ extension is present in the raw assembly. However, read coverage over that region is significantly higher than what is observed over the rest of the mitochondrial genome, thus the two regions most likely exist in different copy numbers and the extension might be the result of missassembly.

**Supplementary Figure 6:**
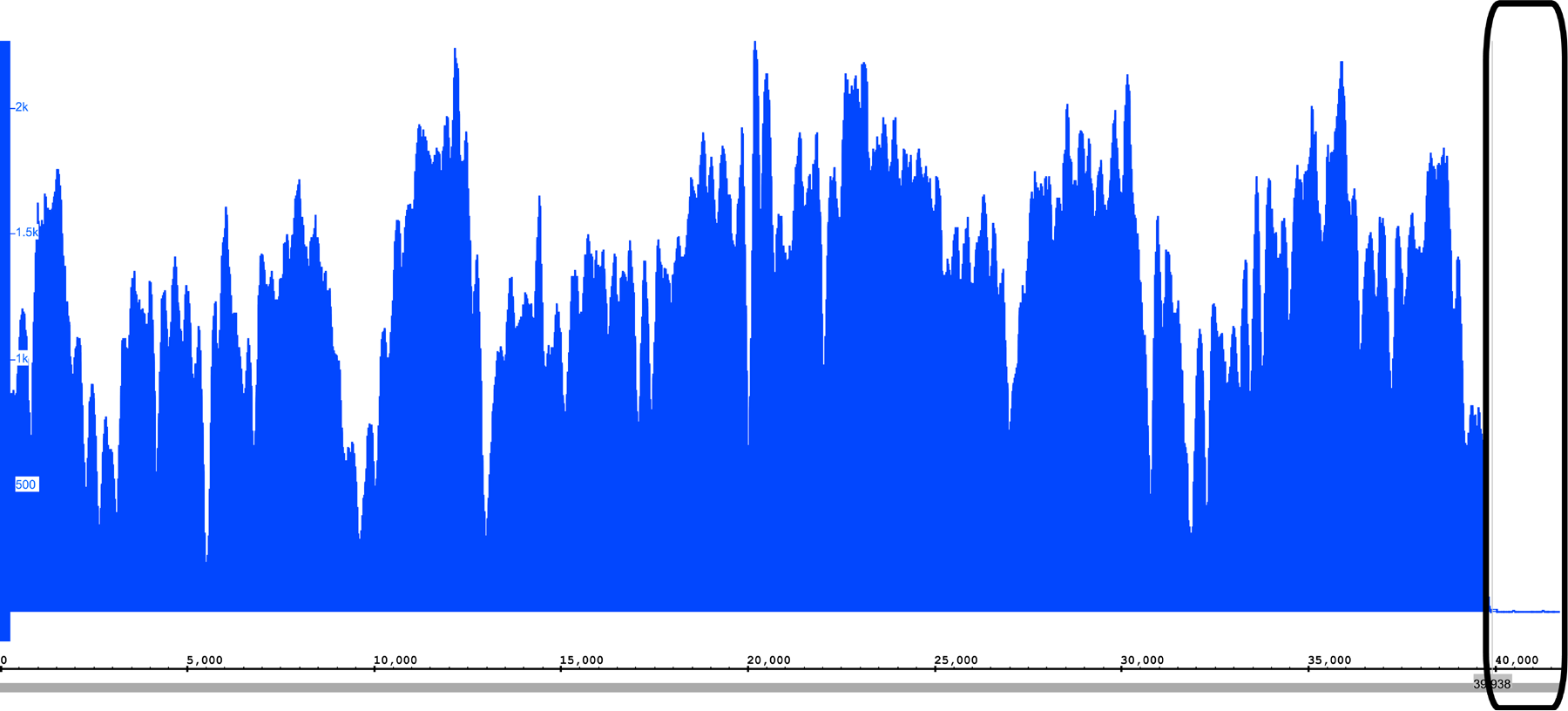
Raw mitogenome assembly and sequencing read coverage for *P. quadecaurelia*. A small 5′ extension is present in the raw assembly. However, read coverage over that region is very low relative to what is observed over the rest of the mitochondrial genome, thus the extension is probably be the result of missassembly.

**Supplementary Figure 7:**
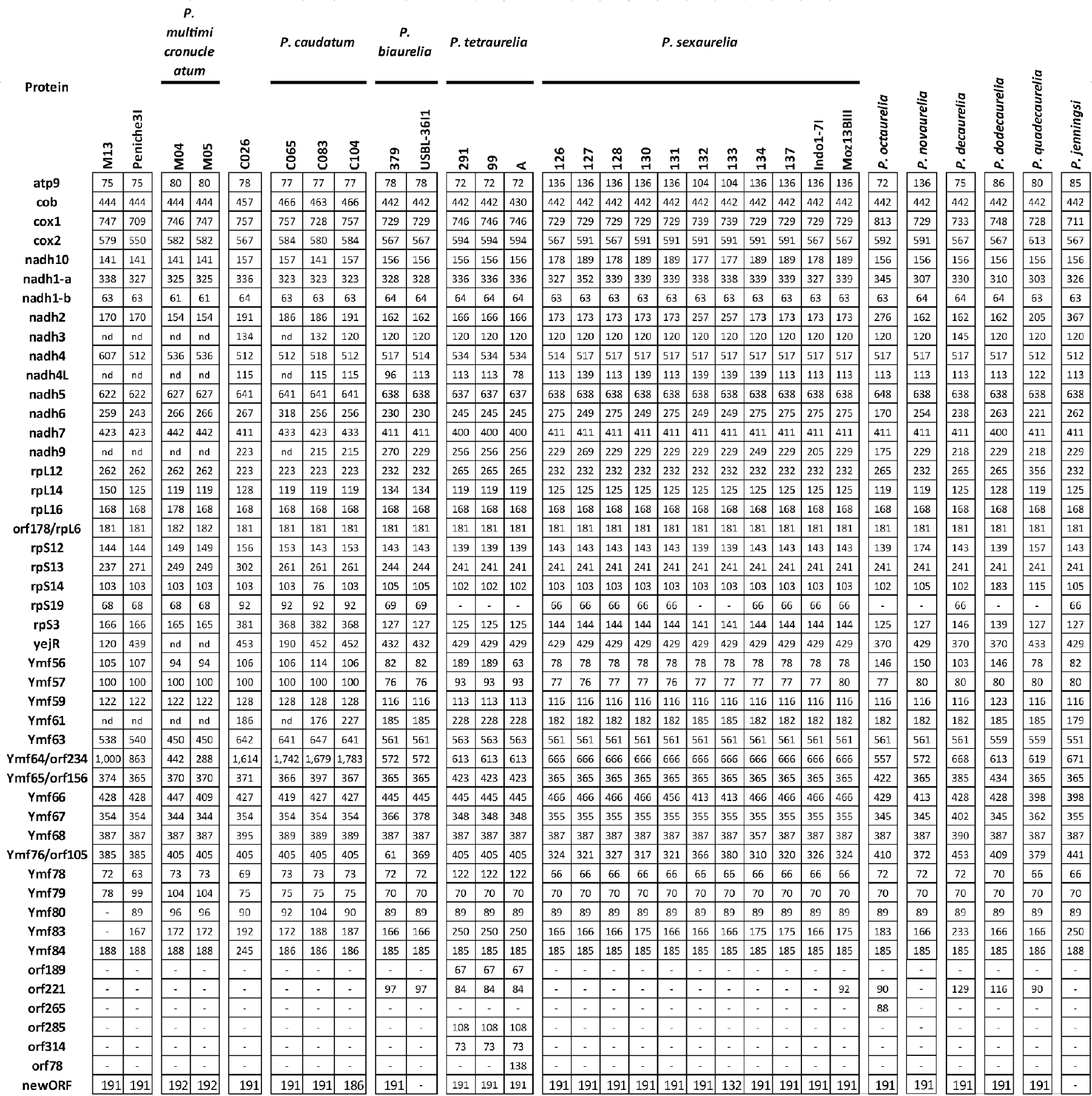
Presence/absence of proteins encoded in mitochondrial genomes in species and individual isolates of species of the *Paramecium* genus. The numbers indicate the length of each protein in amino acids. Most other ORFs exhibit relatively constant length across all *Paramecium* mitochondrial genomes, with the exceptions of *atp9*, which is elongated by ~30-60 AA in all *P. sexaurelia* isolates, *rpS3* and *rps19*, which are specifically elongated in *P. caudatum* and *P. caudatum-C026*, and *Ymf76*, which is very short in one of the *P. biaurelia* isolates. *Ymf64* accounts for a significant part of the difference between the length of *P. aurelia* and *P. caudatum* mitochondrial genomes, being much longer in the latter (~1700 AA vs. ~600 AA). In two isolates of *P. multimicronucleatum* (M04 and M05), the locus appears to have split into multiple ORFs, with the one most clearly homologous to *Ymf64* being shortened even relative to *P. aurelia*. However, in the *P. multimicronucleatum* isolate M13, as well as *P. multimicronucleatum-Peniche3I*, *Ymf64* is relatively long (1000 and 863 AA respectively).

**Supplementary Figure 8:**
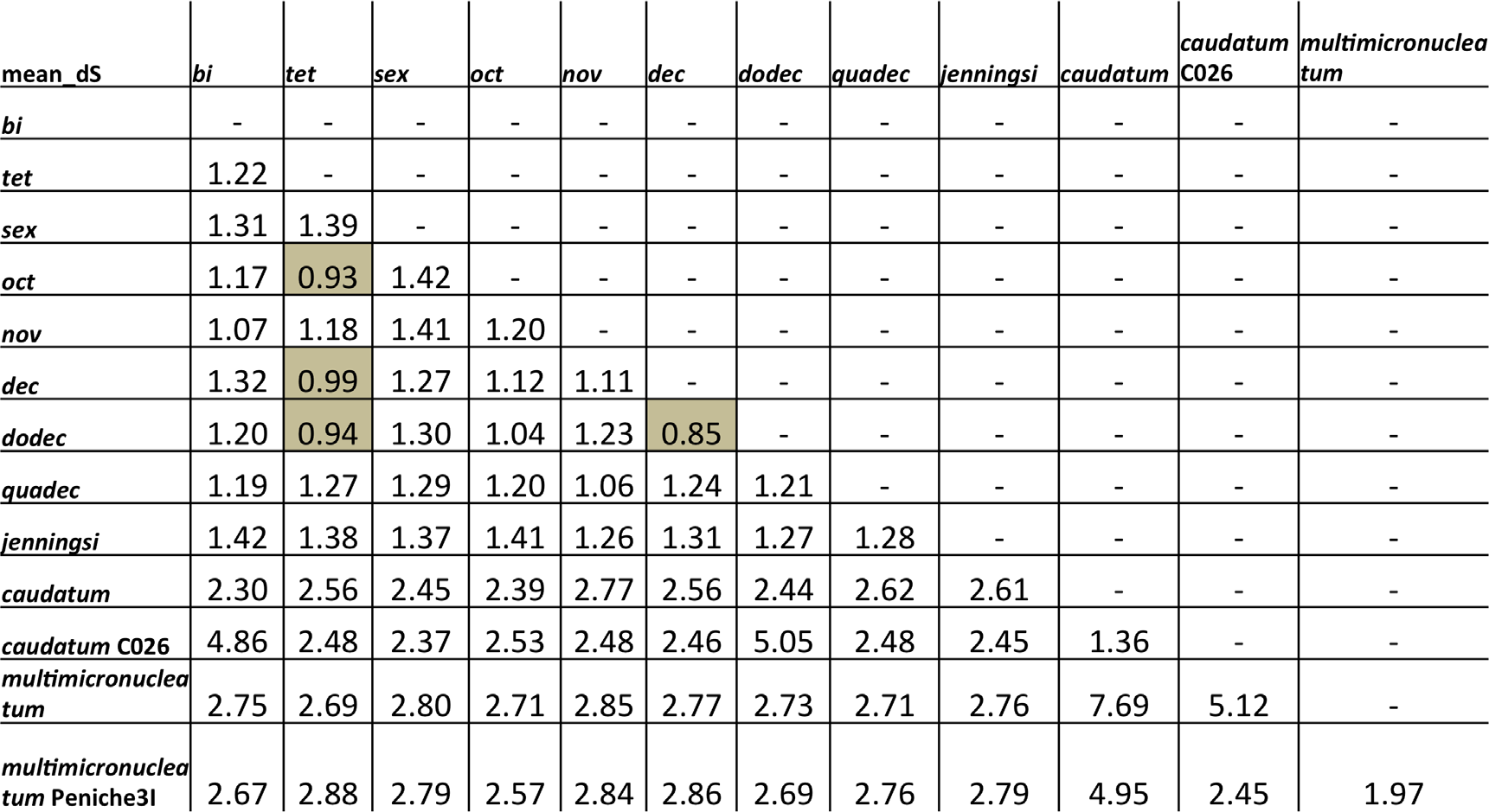
Mean pairwise *dS* between all *Paramecium* species. Species pairs with *dS* < 1.0 are shown in shaded boxes.

**Supplementary Figure 9:**
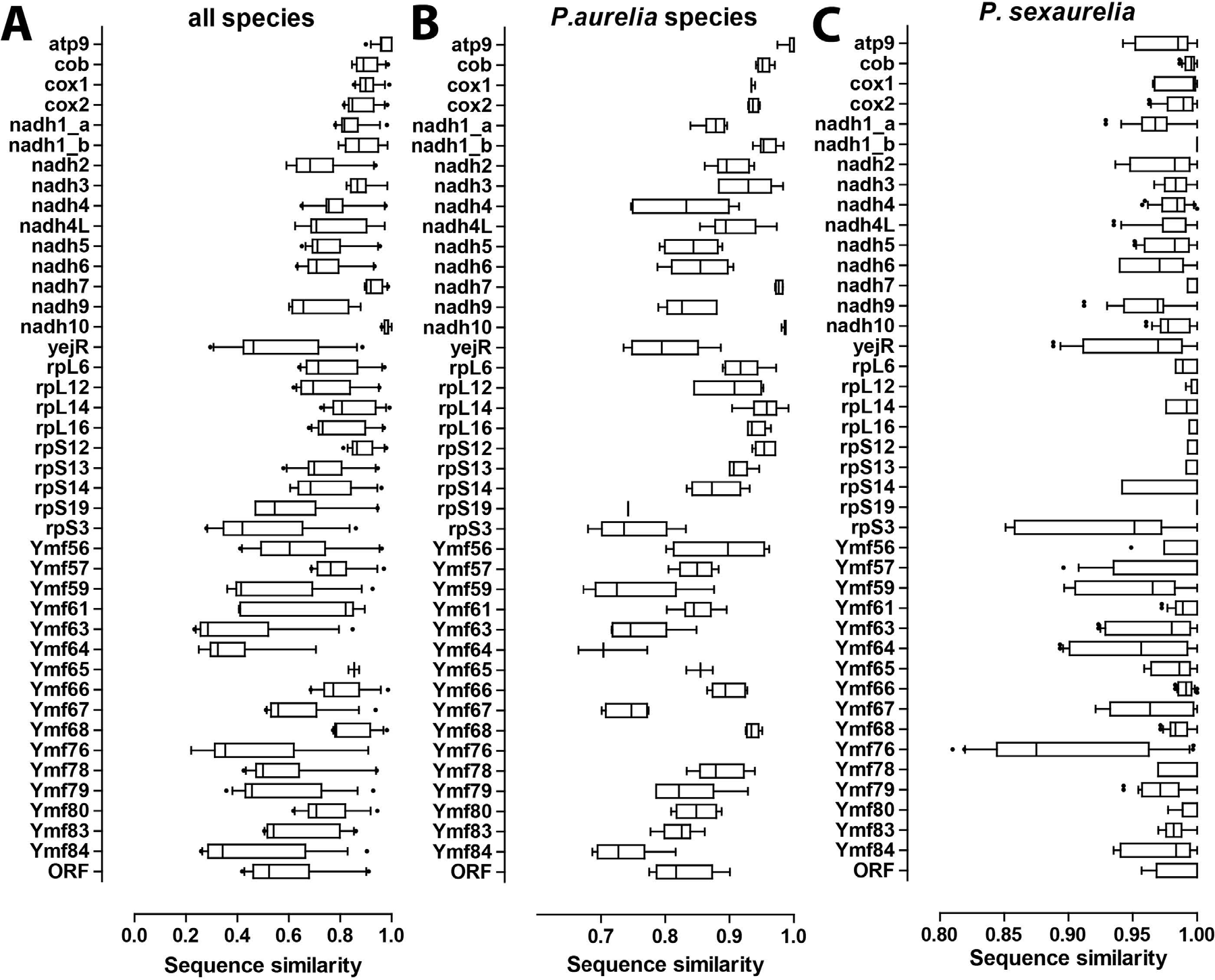
Conservation and divergence of mitochondrial protein sequences in the *Paramecium* genus. Shown is the percent sequence identity for each protein within all species included in this study (A), within the *P. aurelia* species complex (B), and between the 11 individual *P. sexaurelia* isolates (C). Only one isolate per species was included in (A) and (B). The whiskers of the box plot correspond to the 5-95 percentile range.

**Supplementary Figure 10:**
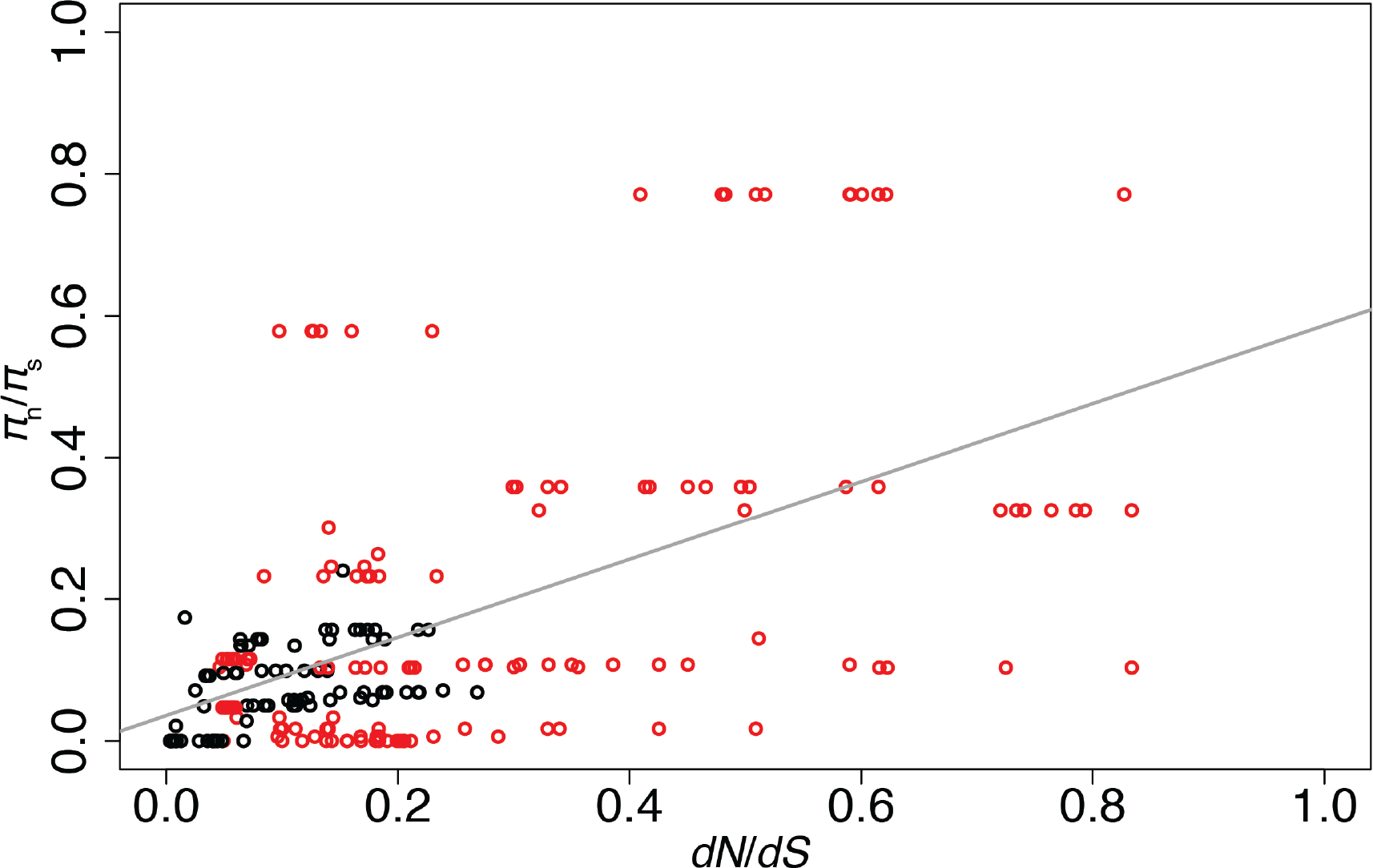
Relationship between observed *π*_n_/*π*_s_ and pairwise *dN/dS* with respect to the closest outgroup species. This analysis was performed for all protein-coding genes, with the conserved genes in black and the *Ymf* genes indicated in red, for *P. tetraurelia, P. sexaurelia, P. caudatum* and *P. multimicronucleatum*. Only genes with *dS* < 1.0 and *θ*_s_ > 1, where *θ*_s_ is the total number of polymorphic sites at synonymous positions, were used in this analysis, resulting in 208 total pairs of observations. This filter was executed in order to reduce errors in *π*_n_/*π*_s_ due to very low values of synonymous polymorphisms in a gene.

**Supplementary Figure 11:**
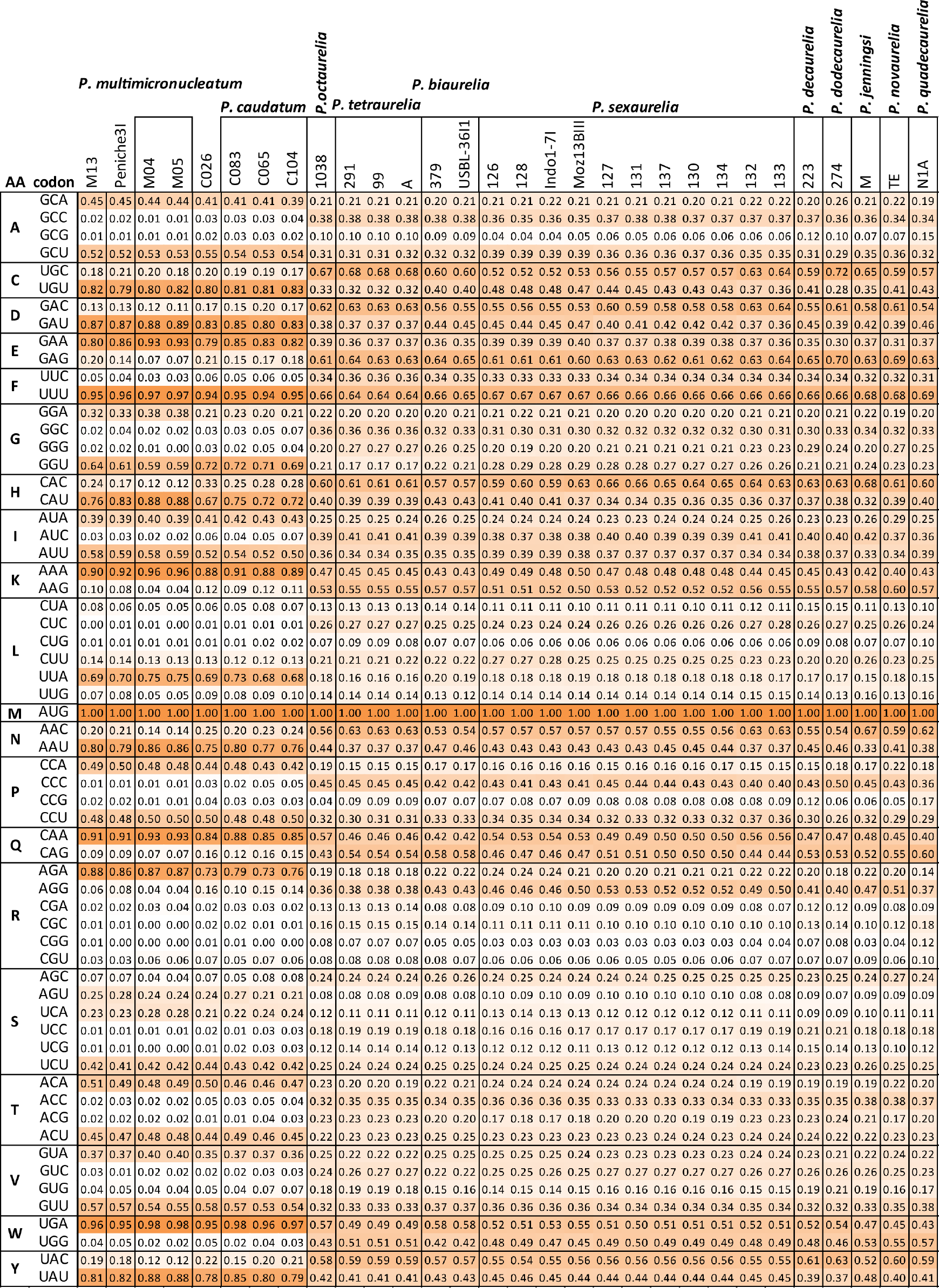
Codon usage in *Paramecium* species. Shown is the frequency of each codon for each amino acid across the Paramecium species included in this study.

**Supplementary Figure 12:**
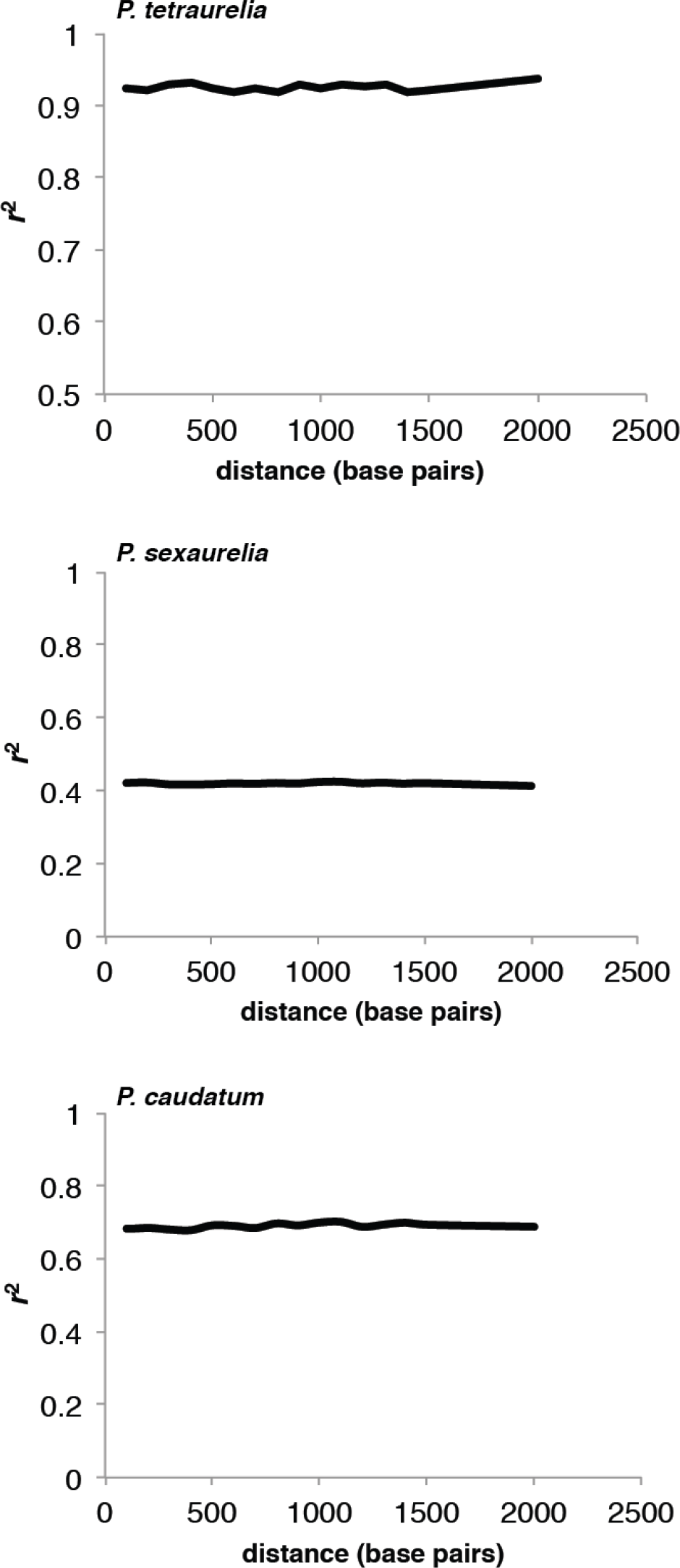
Decay of linkage disequilibrium across the mitochondrial genomes of *P. tetraurelia, P. sexaurelia*, and *P. caudatum*.

**Supplementary Table 1:** *dN/dS* across the *Paramecium* phylogeny (sorted by lowest to highest value of *dN/dS*).

| gene | dN/dS |
| --- | --- |
| NADH dehydrogenase subunit 10 | 0.0071 |
| NADH dehydrogenase subunit 1 b | 0.0159 |
| NADH dehydrogenase subunit 7 | 0.0177 |
| ribosomal protein L14 | 0.0220 |
| apocytochrome_b | 0.0221 |
| Ymf68 | 0.0234 |
| NADH dehydrogenase subunit 3 | 0.0296 |
| NADH dehydrogenase subunit 6 | 0.0305 |
| cytochrome c oxidase subunit 1 | 0.0337 |
| NADH dehydrogenase subunit 4 | 0.0338 |
| Ymf57 | 0.0342 |
| Ymf80 | 0.0347 |
| NADH dehydrogenase subunit 1 a | 0.0396 |
| ribosomal protein S12 | 0.0406 |
| cytochrome c oxidase subunit 2 | 0.0433 |
| ribosomal protein L16 | 0.0482 |
| ribosomal protein L12 | 0.0483 |
| Ymf66 | 0.0491 |
| ribosomal protein L6 | 0.0522 |
| ribosomal protein S13 | 0.0544 |
| NADH dehydrogenase subunit 4L | 0.0559 |
| NADH dehydrogenase subunit 5 | 0.0623 |
| NADH dehydrogenase subunit 2 | 0.0661 |
| Ymf79 | 0.0663 |
| Ymf56 | 0.0703 |
| Ymf78 | 0.0805 |
| ribosomal protein S14 | 0.0818 |
| Ymf65 | 0.0894 |
| Ymf61 | 0.1018 |
| Ymf67 | 0.1105 |
| NADH dehydrogenase subunit 9 | 0.1158 |
| Ymf84 | 0.1200 |
| yejR | 0.1241 |
| Ymf59 | 0.1476 |
| ribosomal protein S3 | 0.1574 |
| Ymf63 | 0.1632 |
| ATP synthase F0 subunit 9 | 0.1781 |
| Ymf83 | 0.1803 |
| Ymf76 | 0.1962 |
| Ymf64 | 0.2154 |

**Supplementary Table 2:** List of genes under positive selection in *P. sexaurelia* and *P. caudatum*. All p values are corrected for multiple tests by Holm Bonferroni method.

| *P. sexaurelia* |  |  |  | *P. caudatum* |  |  |  |
| --- | --- | --- | --- | --- | --- | --- | --- |
| all sites included |  | Using well conserved sites (see methods) |  | all sites included |  | Using well conserved sites (see methods) |  |
| gene | p | gene | p | gene | p | gene | p |
| *Ymf64* | $1.69 \times 10^{-13}$ | *Cox1* | $5.01 \times 10^{-3}$ | *Ymf63* | $5.71 \times 10^{-9}$ | *Ymf64* | $3.48 \times 10^{-7}$ |
| *Ymf67* | $2.71 \times 10^{-7}$ | *Ymf76* | $9.49 \times 10^{-3}$ | *Ymf64* | $2.12 \times 10^{-8}$ | | |
| *Ymf63* | $3.56 \times 10^{-6}$ | *Ymf63* | $2.99 \times 10^{-2}$ | *rpS3* | $2.78 \times 10^{-4}$ | | |
| *Ymf84* | $1.84 \times 10^{-5}$ | | | *Ymf59* | $7.79 \times 10^{-3}$ | | |
| *Cox1* | $2.28 \times 10^{-4}$ | | | *Ymf61* | $1.18 \times 10^{-2}$ | | |
| *Nadh5* | $3.58 \times 10^{-4}$ | | | *Ymf67* | $2.65 \times 10^{-2}$ | | |
| *rpL6* | $1.91 \times 10^{-3}$ | | | | | | |
| *yejR* | $4.22 \times 10^{-3}$ | | | | | | |
| *Ymf65* | $6.72 \times 10^{-3}$ | | | | | | |
| *rpS13* | $8.23 \times 10^{-3}$ | | | | | | |
| *cox2* | $1.05 \times 10^{-2}$ | | | | | | |
| *Ymf76* | $2.72 \times 10^{-2}$ | | | | | | |
| *Ymf61* | $2.79 \times 10^{-2}$ | | | | | | |

**Supplementary Table 3:**
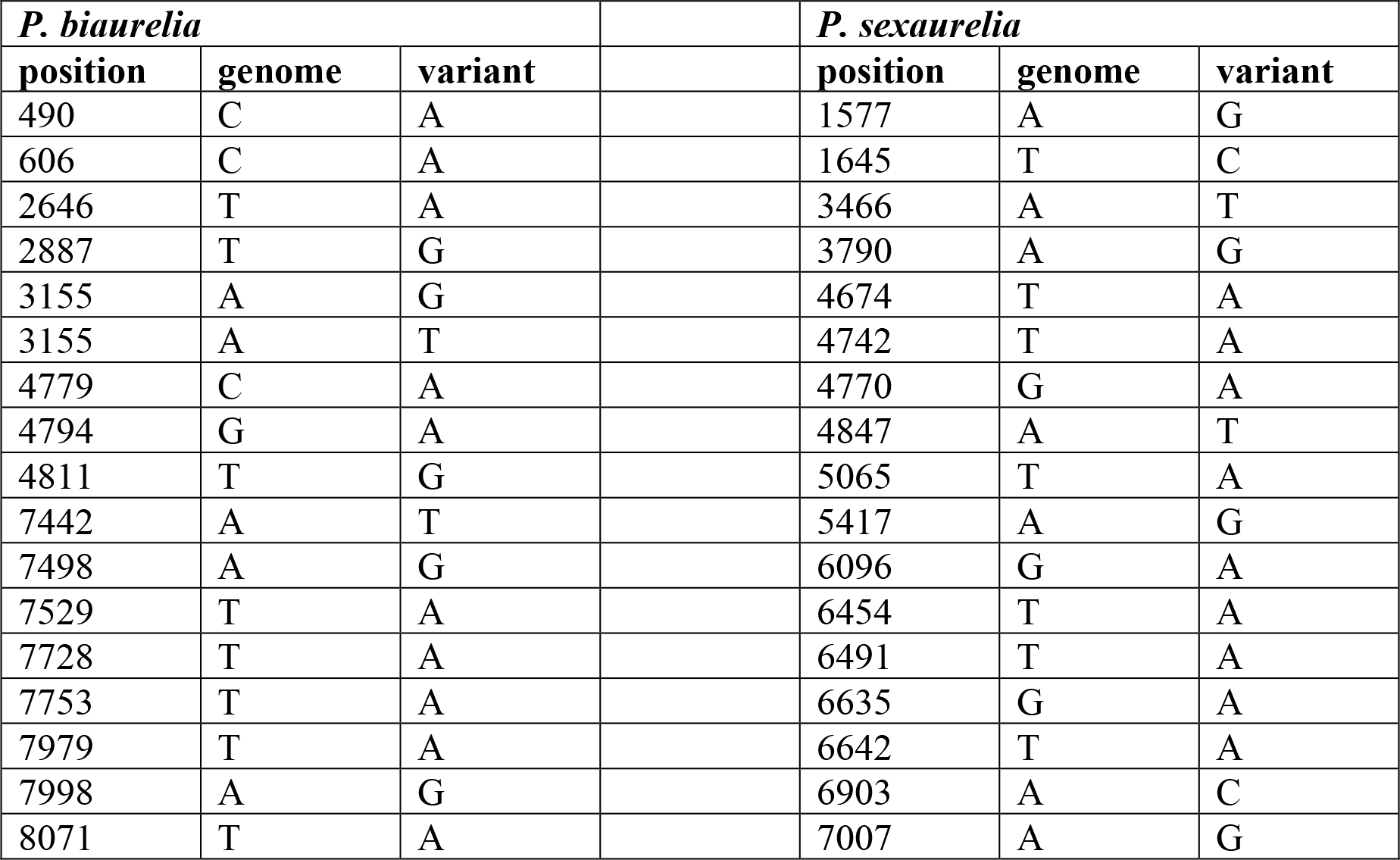
Mitochondrial mutations identified in *P. biaurelia* and *P. sexaurelia* mutation accumulation experiments.

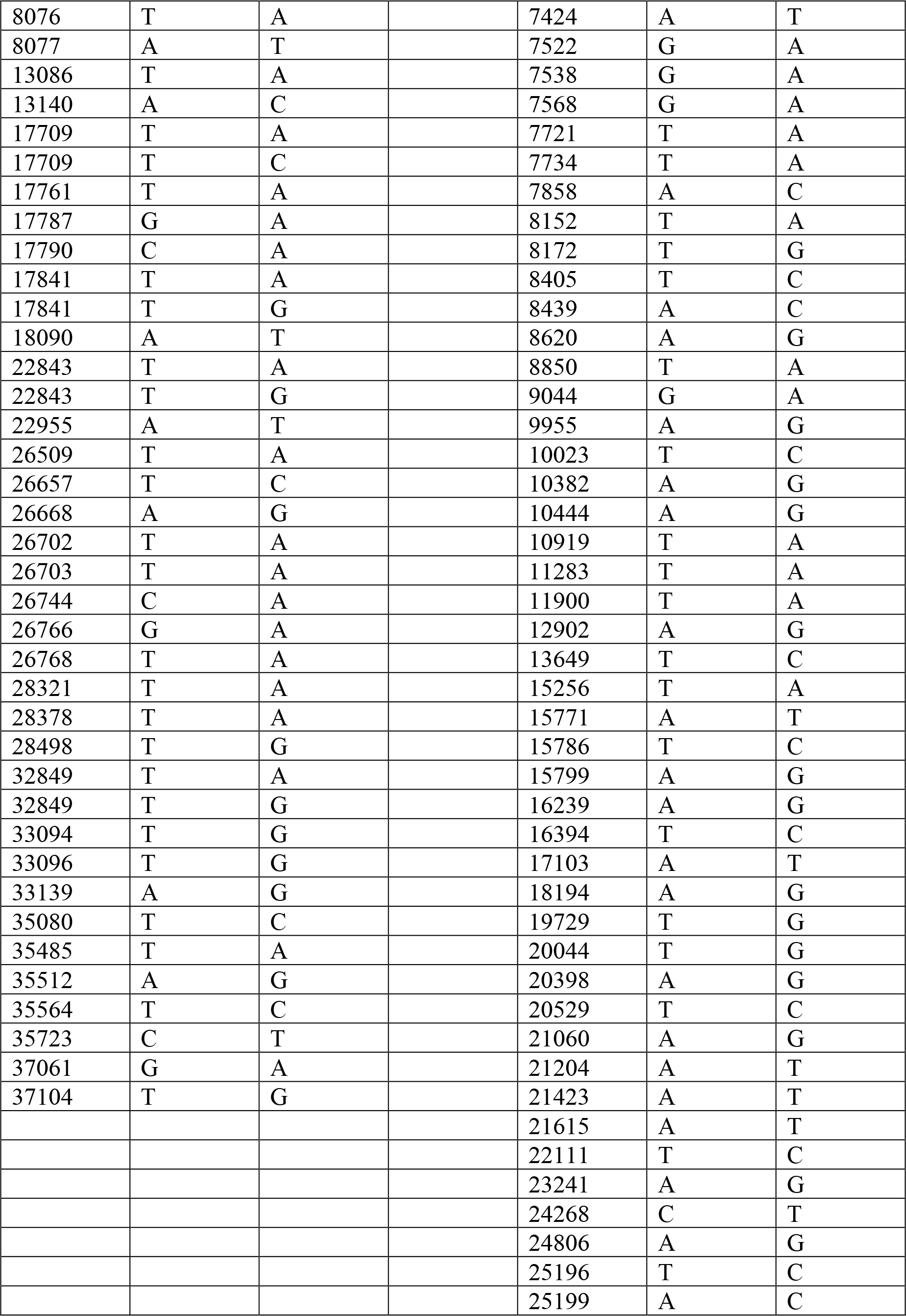

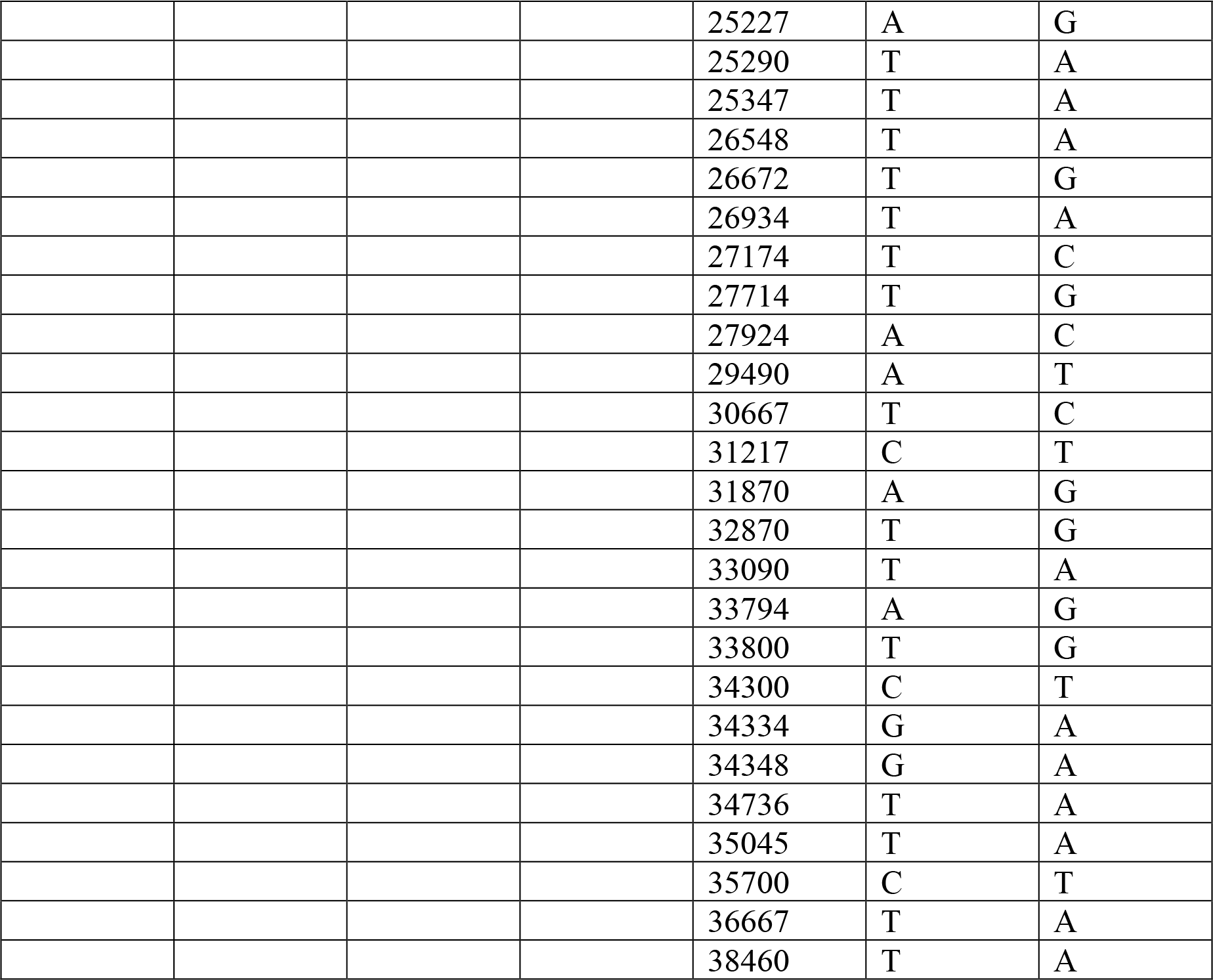

**Supplementary Table 4:**
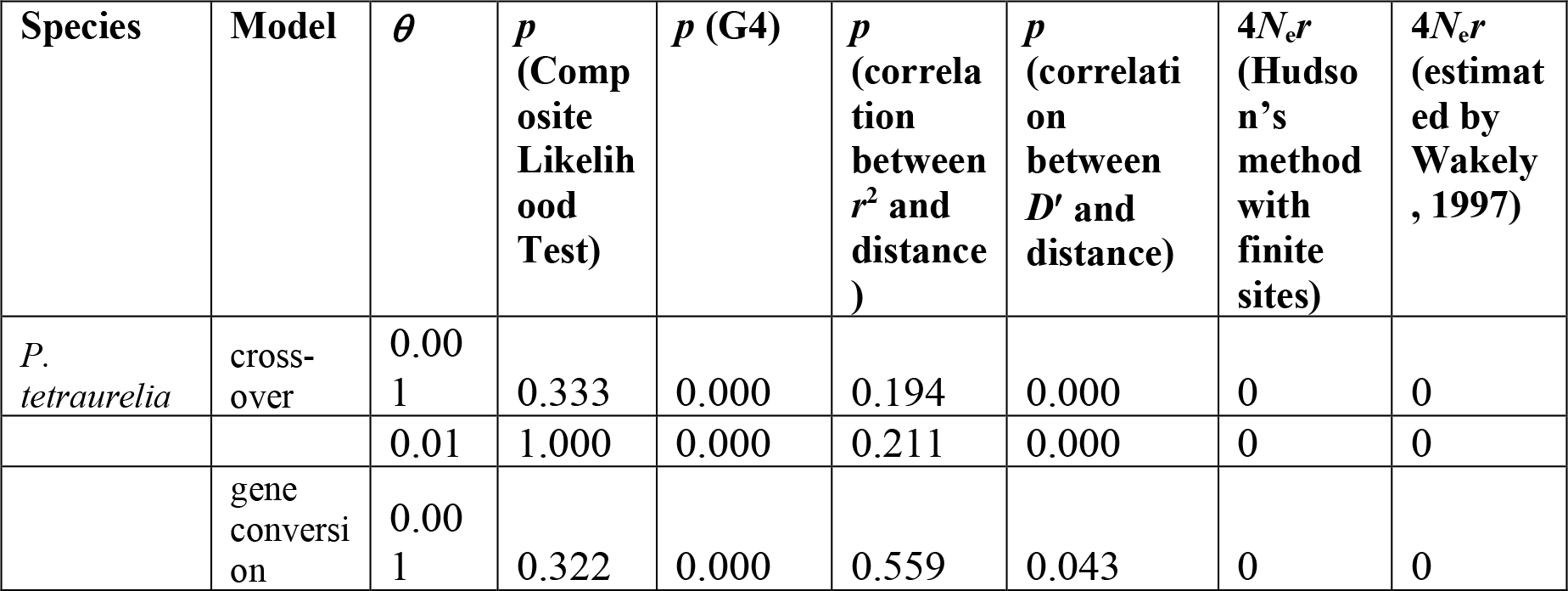
Summary of results obtained from LDhat (pairwise) to detect the presence of recombination in mitochondrial genomes of *Paramecium* species. The *p* values (not corrected for multiple tests) indicate whether values of statistics calculated on observed data are significantly different from the same values obtained for permuted SNPs.

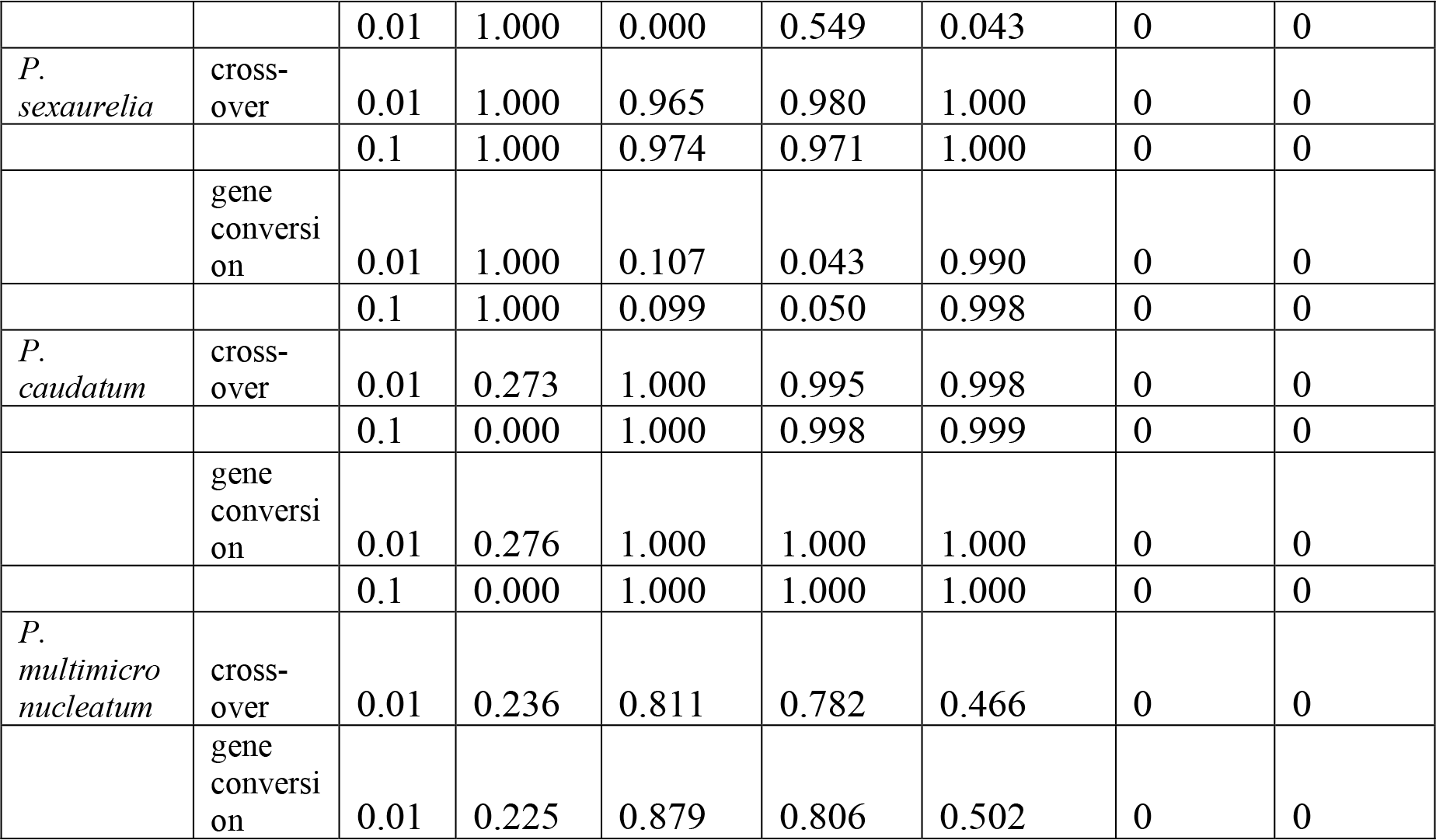

**Supplementary Table 5:** Comparison of neutrality indices between the mitochondrial and nuclear genes that encode for proteins that are part of the oxidative phosphorylation pathway and those that form the structural components of the ribosome.

|  | Oxidative phosphorylation |  |  | Ribosomal proteins |  |  |
| --- | --- | --- | --- | --- | --- | --- |
| | $\pi_n/\pi_s$ | $P_N/P_S/D_N/D_S$ | $\pi_n/\pi_s/dN/dS$ | $\pi_n/\pi_s$ | $P_N/P_S/D_N/D_S$ | $\pi_n/\pi_s/dN/dS$ |
| *P. tetraurelia:* |  |  |  |  |  |  |
| Number of nuclear genes | 27 | 13 | 14 | 336 | 77 | 88 |
| Mitochondrial | 0.127 | 0.781 | 2.384 | 0.062 | 1.935 | 2.248 |
| Nuclear | 0.200 | 2.896 | 6.712 | 0.092 | 0.737 | 1.653 |
| p | 0.223 | 0.285 | 0.422 | 0.387 | 0.352 | 0.262 |
| *P. sexaurelia:* |  |  |  |  |  |  |
| Number of nuclear genes | 22 | 18 | 16 | 429 | 119 | 138 |
| Mitochondrial | 0.124 | 0.328 | 5.180 | 0.035 | 0.089 | 0.754 |
| Nuclear | 0.146 | 1.243 | 7.579 | 0.158 | 1.047 | 3.759 |
| p | 0.535 | 0.444 | 0.145 | 0.093 | 0.124 | 0.308 |
| *P. caudatum:* |  |  |  |  |  |  |
| Number of nuclear genes | 12 | 3 | - | 422 | 9 | - |
| Mitochondrial | 0.105 | 0.364 | - | 0.067 | 0.051 | - |
| Nuclear | 0.133 | 0.211 | - | 0.133 | 0.260 | - |
| p | 0.726 | 0.107 | - | 0.513 | 0.650 | - |

**Supplementary Table 6:** Neutrality indices (defined in the Methods section) in the mitochondria of *Paramecium* species (divergence estimated using all 13 taxa).

|  | *P. tetraurelia* |  | *P. sexaurelia* |  | *P. caudatum* |  |
| --- | --- | --- | --- | --- | --- | --- |
|  | NI<br>[SD] | NI<sub>TG</sub> | NI<br>[SD] | NI<sub>TG</sub> | NI<br>[SD] | NI<sub>TG</sub> |
| All mitochondrial genes | 1.521<br>[2.031] | 0.765 | 0.281<br>[0.475] | 0.213 | 0.500<br>[0.592] | 0.314 |
| Excluding Ymf genes | 1.794<br>[2.515] | 0.690 | 0.339<br>[0.601] | 0.265 | 0.645<br>[0.706] | 0.403 |
| All mitochondrial genes with dS < 1.0 | 1.379<br>[1.454] | 0.829 | 0.539<br>[0.843] | 0.371 | 0.513<br>[0.618] | 0.314 |
| All mitochondrial genes with dS < 1.0, excluding Ymf genes | 1.607<br>[1.890] | 0.715 | 0.859<br>[1.211] | 0.495 | 0.674<br>[0.894] | 0.253 |

